# Small vs. Large Library Docking for Positive Allosteric Modulators of the Calcium Sensing Receptor

**DOI:** 10.1101/2023.12.27.573448

**Authors:** Fangyu Liu, Cheng-Guo Wu, Chia-Ling Tu, Isabella Glenn, Justin Meyerowitz, Anat Levit Kaplan, Jiankun Lyu, Zhiqiang Cheng, Olga O. Tarkhanova, Yurii S. Moroz, John J. Irwin, Wenhan Chang, Brian K. Shoichet, Georgios Skiniotis

## Abstract

Drugs acting as positive allosteric modulators (PAMs) to enhance the activation of the calcium sensing receptor (CaSR) and to suppress parathyroid hormone (PTH) secretion can treat hyperparathyroidism but suffer from side effects including hypocalcemia and arrhythmias. Seeking new CaSR modulators, we docked libraries of 2.7 million and 1.2 billion molecules against transforming pockets in the active-state receptor dimer structure. Consistent with simulations suggesting that docking improves with library size, billion-molecule docking found new PAMs with a hit rate that was 2.7-fold higher than the million-molecule library and with hits up to 37-fold more potent. Structure-based optimization of ligands from both campaigns led to nanomolar leads, one of which was advanced to animal testing. This PAM displays 100-fold the potency of the standard of care, cinacalcet, in *ex vivo* organ assays, and reduces serum PTH levels in mice by up to 80% without the hypocalcemia typical of CaSR drugs. Cryo-EM structures with the new PAMs show that they induce residue rearrangements in the binding pockets and promote CaSR dimer conformations that are closer to the G-protein coupled state compared to established drugs. These findings highlight the promise of large library docking for therapeutic leads, especially when combined with experimental structure determination and mechanism.

**One sentence summary:** Structure-based virtual screening uncovers novel CaSR allosteric modulators with enhanced efficacy and less side effects.

## Introduction

Well before the advent of molecular pharmacology, much effort had been directed toward developing “calcimimetic” and “calcilytic” drugs to promote or suppress the calcium-sensing abilities of parathyroid cells and to regulate PTH secretion and blood calcium levels. The activity of these drugs on the calcium-sensing receptor (CaSR), a G protein-coupled receptor (GPCR), was confirmed after its cloning (*1*). CaSR is present in almost every organ system but is most highly expressed in the parathyroid gland and in the kidneys, where it maintains calcium homeostasis by sensing changes in extracellular calcium levels to regulate PTH secretion, renal calcium reabsorption, and excretion (*2, 3*). Loss-of-function mutations or reduced CaSR expression cause familial hypocalciuric hypercalcemia (FHH), neonatal severe primary hyperparathyroidism, or adult primary hyperparathyroidism, respectively (*4*). In FHH, the CaSR becomes less sensitive to rising calcium levels, leading to increased PTH secretion *in lieu* of elevated blood calcium levels and reduced calcium excretion. Conversely, oversensitivity to calcium from gain-of-function mutations in autosomal dominant hypocalcemia (ADH) decreases PTH secretion and lowers blood calcium levels (*5–7*). Through its widespread expression, CaSR is also involved in other physiological mechanisms, notably gastrointestinal nutrient sensing, vascular tone, and secretion of insulin, with alterations in receptor activity implicated in the development of osteoporosis and in several cancers (*3*).

Efforts to target CaSR therapeutically have focused on the development of positive and negative allosteric modulators (PAMs and NAMs), which potentiate the receptor’s activation or its inactivation, respectively, while binding at a non-orthosteric site (here, a non-calcium site). PAMs enhance the physiological response to calcium but display little or no agonist activity on their own. In the past two decades, the small molecule PAM drug cinacalcet and the peptide-based PAM drug etelcalcetide (*8*) were approved for human use, but only for the treatment of secondary hyperparathyroidism (HPT) in patients with chronic kidney disease (CKD) undergoing dialysis (usually stage 5), while cinacalcet is also approved to treat high levels of calcium in patients with parathyroid cancer. The limited indications reflect the adverse side effects associated with the current PAMs, including hypocalcemia, gastrointestinal problems, hypotension, and adynamic bone disease (*9*). Hypocalcemia is life-threatening as it can cause seizures and heart failure (*10–15*). CKD affects more than 10% people worldwide and considering the prevalence of secondary hyperparathyroidism in various stages of (*9, 16–20*), drugs that decrease PTH levels without causing hypocalcemia are much needed.

The CaSR belongs to the Family C of GPCRs, a relatively exotic group of receptors that have the unique property of operating as homo- or heterodimers with extracellular domains (ECDs) constituting the orthosteric ligand binding site. The ECD of a CaSR monomer is connected through a linker region to the seven transmembrane domain (7TM), which has been shown to activate primarily Gq/11 and Gi/o G protein subtypes to elicit signaling (*21, 22*). Upon calcium binding to the ECDs, the CaSR homodimer undergoes extensive conformational transitions that bring the 7TMs in close proximity through a TM6-TM6 interface, an overall configuration that has been shown to be associated with receptor coupling to G protein (*23, 24*). Our recent high resolution cryo-EM studies showed that in the active-state receptor both cinacalcet and the related evocalcet, recently approved for therapeutic use in Japan (*22*), both adopt an “extended” conformation within the 7TM of one CaSR monomer, and a “bent” conformation in the second monomer of the dimer. The two different conformations by the same ligand reflect changes in the allosteric PAM binding pockets that are transforming to accommodate the asymmetric juxtaposition of the two CaSR protomers upon activation (*22*).

We sought to exploit these structures and our mechanistic insights on receptor activation to discover new CaSR PAM chemotypes that are topologically unrelated to those previously investigated. Such new chemotypes often lead to new pharmacology, and our hope was that they might enhance CaSR activation and so modulate PTH secretion without leading to the dose-limiting hypocalcemic actions of approved drugs. To address this, we adopted a structure-based, virtual library docking approach (*25*). In the last four years, docking libraries have expanded over 1000-fold, from millions to billions of molecules, and from these new libraries have emerged unusually potent ligands, with activities often in the mid- to low-nM concentration range, straight from docking (*25–31*). Indeed, simulations suggest that as the libraries expand, docking finds not only more but better ligands, although this has not been experimentally tested. While our chief goal was the discovery of efficacious CaSR PAMs with reduced side-effects, we took the opportunity to test how library growth affected docking experimentally, comparing the *in vitro* results from docking a 2.7 million library vs. a library of 1.2 billion molecules. This offers one of the first experimental tests for the impact of library growth on experimental outcome. Mechanistically and therapeutically, potent new PAMs emerged from these studies, active in the 3 nM range, with *in vivo* activities between 10 and 100-fold more potent than cinacalcet, and apparently without that drug’s dose-limiting hypocalcemia. Cryo-EM structures of the new PAMs illuminate their mechanism of action on CaSR and may template future optimization and discovery toward better therapeutics.

## Results

### Docking in-stock and ultra-large make-on-demand library against CaSR for new PAMs

We began by docking the smaller, in-stock library of 2.7 million molecules at both 7TM sites of CaSR (**Fig. 1A, fig. S1A**). In the site accommodating the “bent” conformation of cinacalcet (7TM^B^ site), an average of 3,927 orientations of each library ligand were sampled, each in an average of 333 conformations, or 1.2 trillion configurations overall; the calculation took just under one hour of elapsed time on a 1000-core cluster, using DOCK3.7 (*32*). Molecules were scored for van der Waals (*33*) and Poisson-Boltzmann-based electrostatic complementarity (*34, 35*) corrected for Generalized-Born ligand desolvation (*36*). Conformationally strained molecules were deprioritized (*37*), while high-ranking molecules were clustered for similarity to each other using an ECFP4-based Tanimoto coefficient (Tc) of 0.5 and filtered against similarity to known CaSR ligands. Comparable numbers of ligand orientations, conformations, and docking configurations were sampled and calculated for the “extended” site (7TM^A^ site). Ultimately, we selected 26 compounds with favorable interactions at the 7TM^A^ site, and 22 compounds with interactions at the 7TM^B^ site (**fig. S1A**). These were tested for CaSR-induced G_i3_ activation (*38*) using an extracellular calcium concentration of 0.5 mM. One PAM emerged from those selected for the 7TM^A^ site with > 10% of *Emax* induced by cinacalcet, and three PAMs were found for the 7TM^B^ site, representing hit rates of 3.8% (1/26) and 13.6% (3/22), respectively (**fig. S1A**). The higher hit rate for the 7TM^B^ site is likely attributed to its more enclosed pocket, which better excluded molecules unlikely to bind and led to better ligand complementarity.

**Fig. 1.**
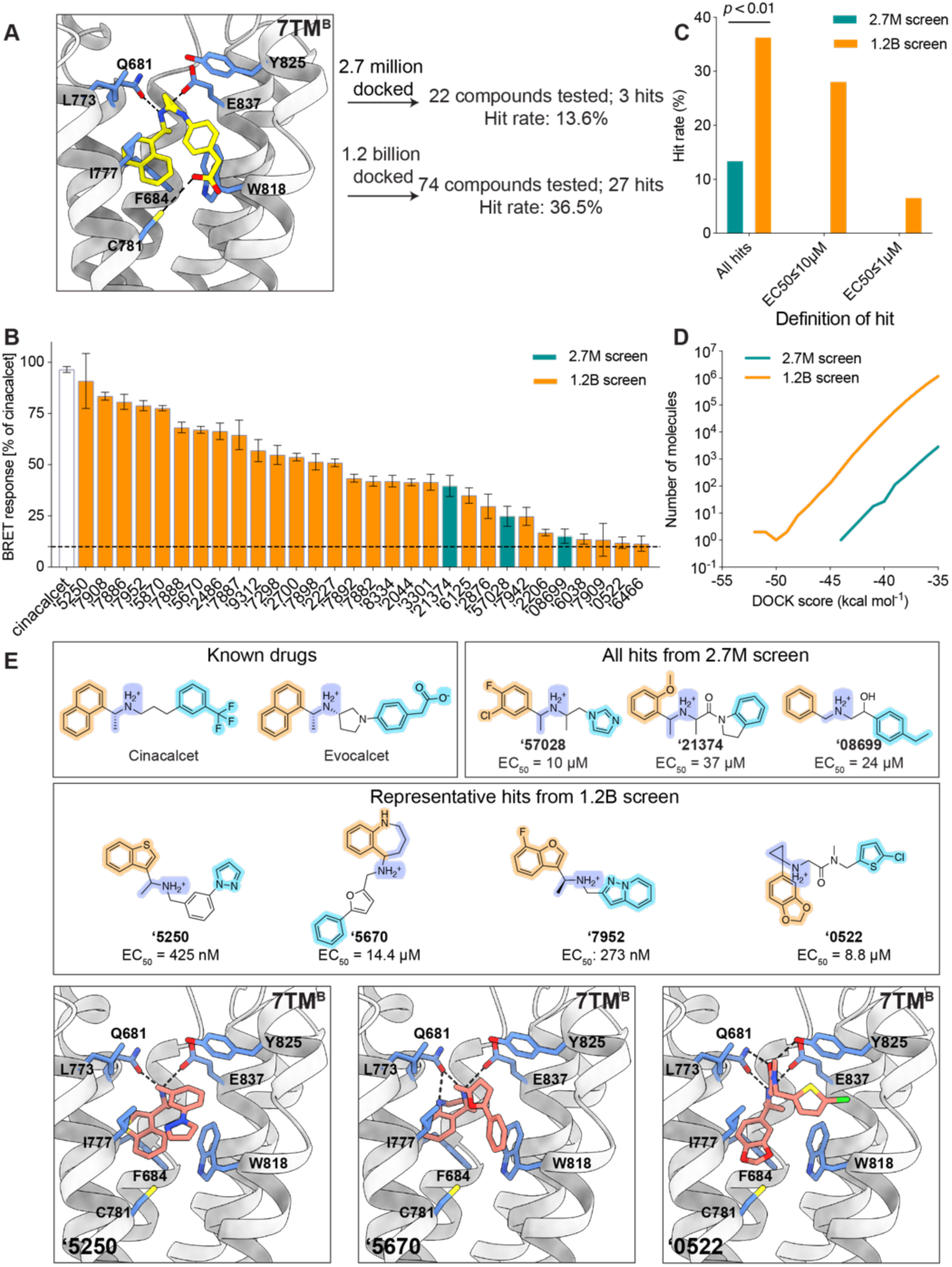
Novel ligands identified from the in-stock and large library screens targeting the 7TM sites of CaSR. (**A**) Larger scale docking against the 7TM^B^ site results in higher hit rate (13.6% in 2.7-million docking campaign versus 36.5% in 1.2-billion docking campaign). Hit rates were defined by over 10% BRET response compared to cinacalcet at 100 μM. E_VDW_: van der Waals; E_ES_: electrostatic; E_LDS_: ligand desolvation. Cinacalcet is in gold and evocalcet is in pink for illustration of the binding sites (**PDB: 7MCF**). (**B**) BRET response (normalized to cinacalcet) of the initial hits at 100 μM. (**C**) Hit rate comparison between 2.7 million and 1.2 billion screens with different affinity definitions. The overall hit rate of the 1.2 billion screen is significantly better than the in-stock 2.7 million screen (*P* < 0.01 by z-test). (**D**) Total docking energies of top-scoring molecules out of LSD compared to in-stock screen (only molecules with DOCK score < -35 kcal mol^-1^ are plotted). (**E**) Examples of the docking hits in comparison to the known PAM drugs cinacalcet and evocalcet (colors represent the different moieties fulfilling the same role). Docked poses of the novel PAM representatives at two 7TM sites are shown.

To measure the impact of larger libraries, and potentially identify more potent PAMs, we screened a library of 1.2 billion make-on-demand (“tangible”) molecules (*39*) against the more enclosed 7TM^B^ site (**Fig. 1A**). Here, an average of 1,706 orientations were sampled for each library molecule, each in an average of 425 conformations, or 682 trillion total configurations overall; this calculation took about 16 days of elapsed time on a 1,000-core cluster. Top-scoring molecules were filtered and clustered as for the smaller library, and 1,002 cluster-heads passed all criteria. 96 molecules that score well in both sites were prioritized for synthesis, of which 74 compounds were successfully made, a 77% fulfillment rate (**Fig. 1A, fig. S1B**). In BRET assays, 27 of the 74 compounds produced >10% of the *Emax* induced by cinacalcet, a 36.5% hit rate and almost three-fold higher than did the 2.7 million molecule library (**Fig. 1, B to C**).

The larger library also revealed hit molecules with higher potency than those from the smaller library, with more than 70% having EC_50_ values better than 10 µM, and 20% having an EC_50_ better than 1 µM (**Fig. 1C, table S1**), with the best having an EC_50_ of 270 nM. Given the number of molecules tested, the hit-rate difference between the larger and smaller library screens was significant (*p*-value < 0.01), and in fact is only as good as it is for the smaller library when we count as hits molecules with EC_50_ values worse than 10 µM. In the 1-10 µM and in the 0.1-1 µM ranges, no hits emerged from the smaller library, whereas multiple ones did so from the larger library. These results support the theoretical studies predicting better performance from larger libraries (*40*), providing an experimental quantification for the impact of larger versus smaller libraries (**Fig. 1, C to D**). The ultra-large library also demonstrates superior performance in terms of chemical novelty. While the compounds identified from the in-stock screens were topologically distinct from known ligands in the ChEMBL and IUPHAR databases, with ECFP4 Tanimoto coefficients (*Tc*) less than 0.35 as shown in **table S1**, they still exhibited physical similarities to established PAMs. These similarities include a buried aromatic ring, a bridging methylamine linker, and a distal aromatic ring, as illustrated in **Fig. 1E**. In contrast, the PAMs identified from the ultra-large library showcased a diverse range of heteroaromatic anchors, various linker types, or even the absence of a linker as in the case of compound **‘7909** (**table S1, fig. S1B**). Notably, many of these compounds, such as **‘5670** and **‘0522**, lacked the methyl group adjacent to the cation, a feature commonly found in CaSR PAMs (**Fig. 1E**).

Our recent cryo-EM structures have shown that the cationic amine of cinacalcet and evocalcet hydrogen bonds and ion-pairs with Q681^3.33^ and E837^7.32^ of CaSR, which are critical for PAM recognition (*22, 41*). Meanwhile, the highly conserved methyl group α to this cationic amine fits into a hydrophobic pocket formed by I841^7.36^, F684^3.36^, F668^2.56^, whose substitutions with alanine abolish or decrease binding affinities for CaSR PAMs (*42*). In their bent conformations bound to 7TM^B^, the naphthalene ring common to both drugs T-stacks with F684^3.36^ and W818^6.50^, while the benzene forms edge-to-pi interaction with W818^6.50^. Remarkably, although their linker lengths differ from the known drugs, most of the new PAMs also adopt “bent” conformations in their docked poses within the 7TM^B^ pocket (**Fig. 1E, fig. S1**). While most of them retain hydrophobic flanking groups that dock into the aryl sites defined by cinacalcet and evocalcet, all do so with different moieties (**Fig. 1E**, **Fig. 2A**).

**Fig. 2.**
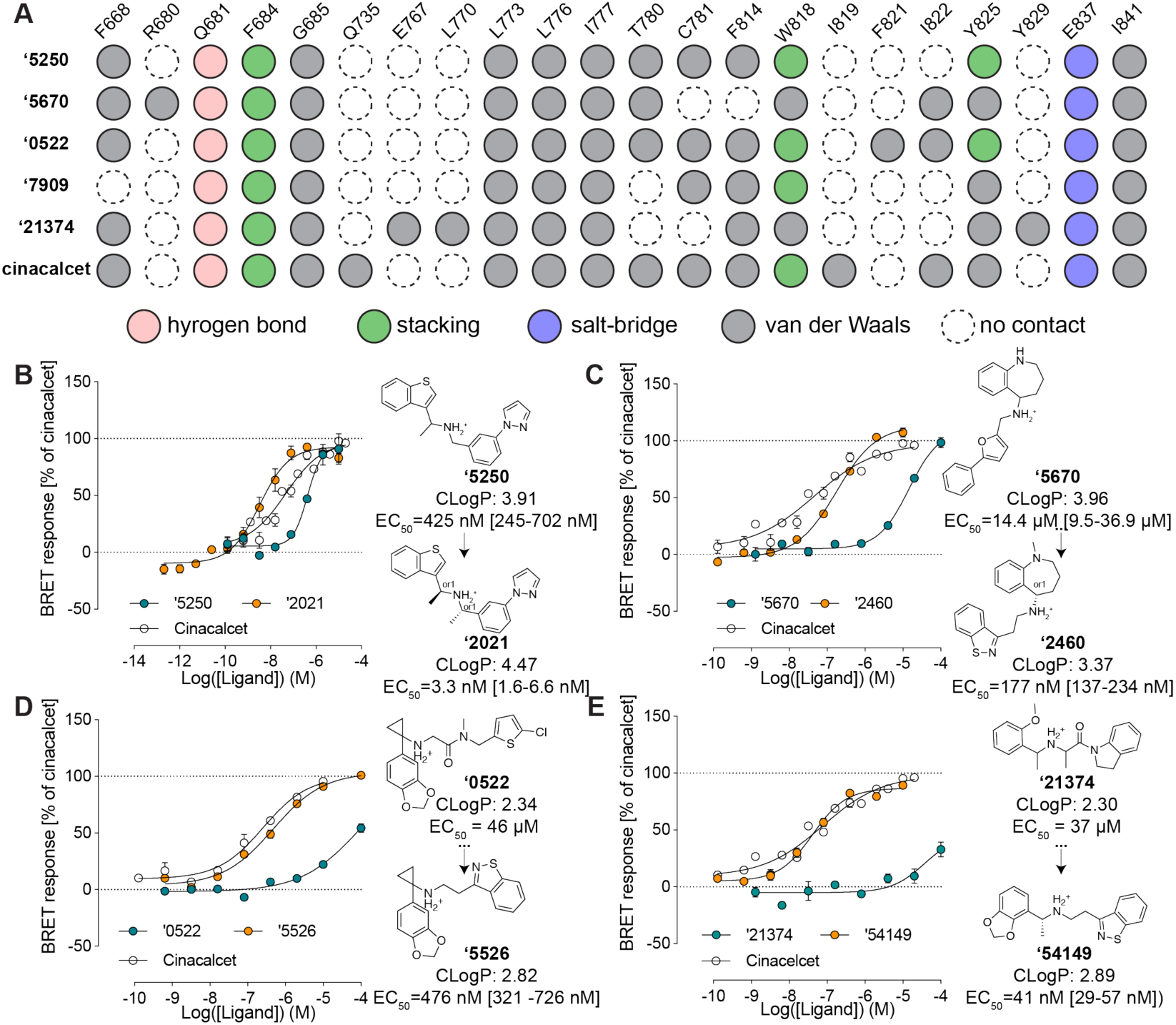
Initial hits to high-affinity analogs. (**A**) Contact analyses of the initial docking hits versus cinacalcet. (**B**) Docking hit **‘5250** and its optimized analog **‘2021** (a diastereomer of **‘6783**). (**C**) Docking hit **‘5670** and its optimized analog **‘2460** (an enantiomer of ‘6218). (**D**) Docking hit **‘7909** and its optimized analog **‘3161**. (**E**) In-stock docking hit **‘21374** and its optimized analog **‘54149**. EC_50_ was determined by monitoring Gi activation by CaSR upon compound addition at [Ca^2+^] = 0.5 mM. The efficacy of the compounds is normalized to the maximum BRET response induced by cinacalcet. Data represent means and SEMs of 3-27 replicates.

### Structure-based optimization of new PAMs

A core goal of this study was finding new chemotypes conferring new pharmacology. We therefore prioritized high-ranking docked molecules based on both potency and topological dissimilarity to known CaSR PAMs for further optimizations. To increase the affinity of the initial hits, we sought to optimize interactions with residues that had proven important in other series^33^, including F668^2.56^, Q681^3.33^, E837^7.32^, I841^7.36^, F684^3.36^, and W818^6.50^ (**Fig. 2A**). The greater polarity of the docking hits, whose calculated octanol:water partition coefficient (clogP) ranged from 2.3 to 4.0 vs. a clogP of 5.1 for the more hydrophobic cinacalcet, gave us freedom to operate in the hydrophobic CaSR site.

To fill a gap in the interface with L773^5.40^, Y825^6.57^ and to stiffen the linker in the docking-derived PAM **‘5250** (EC_50_ 415 nM), a second methyl was added proximal to the cationic nitrogen. This improved potency five-fold, to an EC_50_ of 90 nM, while synthetic resolution of the diastereomers improved it another 130-fold. The resulting compound Z8554052021 (**‘2021**), with an EC_50_ of 3.3 nM, is among the most potent CaSR and indeed GPCR PAMs ever described (**Fig. 2B**).

The tetrahydrobenzapine of compound ‘**5670** (**Fig. 2C**) separates it from the naphthalene equivalent of cinacalcet and evocalcet and gives it a relatively polar and certainly three-dimensional character compared to the equivalent groups of other CaSR PAMs. Substitution of the terminal phenyl-furan with a more compact and more polar benzothiazole, which can be well-accommodated in the hydrophobic site defined by residues I777^5.44^, W818^6.50^ and Y825^6.57^, improved potency seven-fold (compound Z2592185946 (**‘5946**)), while its N-methylation led to **‘6218** (EC_50_ 0.25 µM) (**fig. S3B**). Enantiomeric purification led to **‘2460**, a 177 nM CaSR PAM (**Fig. 3C**). Despite its 80-fold potency improvement, the molecular weight and cLogP values of **‘2460** were actually reduced versus the parental docking hit, improving Lipophilic Ligand Efficiency (LLE) from 0.9 to 3.4. Furthermore, substituting the nitrogen atom in tetrahydrobenzapine with oxygen, sulfur, or carbon results in the inactivation of the compounds, thereby making them ideal probe pairs for physiological studies (**fig. S3B**).

**Fig. 3.**
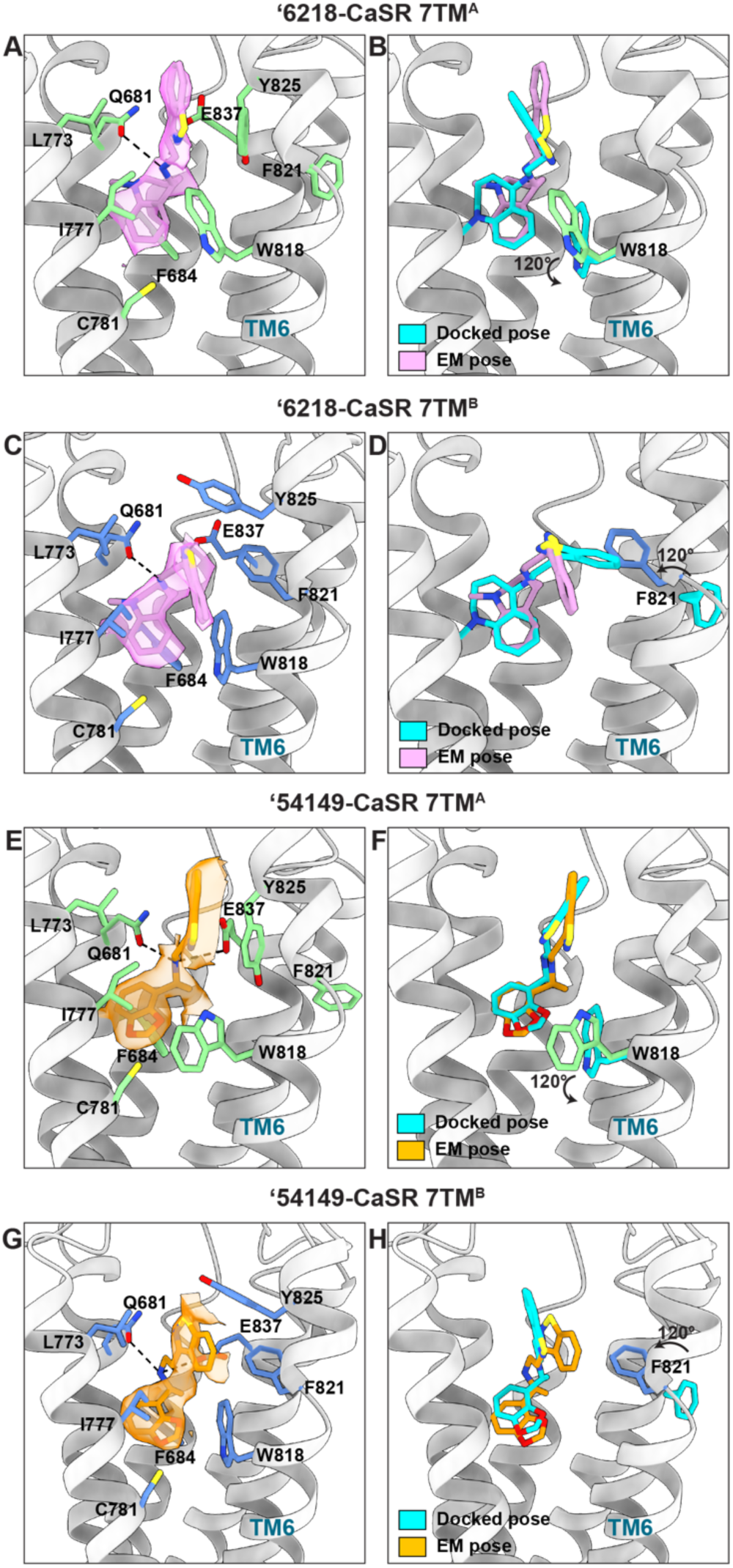
Structural comparison between docked and experimentally determined poses for ‘54149 and ‘6218. (**A**) Close-up view of **‘6218** in the 7TM^A^ site, with its EM density shown. Surrounding residues are in green. (**B**) Superposition of docked and experimentally determined pose of **‘6218** in the 7TM^A^ site. (**C**) Close-up view of **‘6218** in the 7TM^B^ site, with its EM density. Surrounding residues are shown in blue. The docked pose and its surrounding residues are in silver. (**D**) Superposition of docked and experimentally determined pose of **‘6218** in the 7TM^B^ site. (**E**) Close-up view of **‘54149** in the 7TM^A^ site, with its EM density. The surrounding residues are in green. (**F**) Superposition of docked and experimentally determined pose of **‘54149** in the 7TM^A^ site. (**G**) Close-up view of **‘54149** in the 7TM^B^ site, with its EM density. The surrounding residues are in blue. (**H**) Superposition of docked and experimentally determined pose of **‘54149** in the 7TM^B^ site. (**B**, **D**, **F**, **H**) The residues undergoing conformational changes in the experimental structures are shown. Docked poses and protein residues in the docked structures are in cyan.

Similar changes in the equivalent aryl groups, binding in the hydrophobic site defined by residues F668^2.56^ and I841^7.36^, led to improvements in docking hits Z5208267909 (**‘7909**) and Z1591490522 (**‘0522**) (**Fig. 2D, fig. S3D**). For the former, the EC_50_ improved from over 100 µM to 1.7 µM (Z6562953161 (**‘3161**)**, fig. S3D**), and efficacy was much improved even though molecular weight was, again, decreased. Meanwhile, the analog of **‘0522**, Z6923555526 (**‘5526**), saw the introduction of the same benzothiazole as in ‘**2460**, along with a simplification of the linker, giving better complementarity with the hydrophobic site defined by I777^5.44^, W818^6.50^ and Y825^6.57^ (**fig. S3C**), and improving EC_50_ 95-fold, to 0.48 µM.

We also sought to optimize the early PAMs revealed by the “in-stock” library. Although these molecules began with weak EC_50_ values, we were able to optimize three of the four molecules to EC_50_ values of 30-163 nM by 2D searching through a 46 billion make-on-demand catalog library in SmallWorld (https://sw.docking.org/) (**Fig. 3E, fig. S2**). Most compelling was the improvement of **‘21374**. Here, simplification of the linker and installation of a benzothiazole, as in **‘6218** and **‘5526**, above, led to **‘85339**, with an EC_50_ of 174 nM. Stereochemical purification to (*R)-***‘85339** (**‘54149**) revealed a 41 nM PAM. It was this molecule, with relatively high potency (4.5-fold improved on that of cinacalcet), favorable cLogP (2.9), new chemotype, and novel receptor contacts, that we ultimately took forward into *in vivo* studies.

### Cryo-EM structures of the ‘6218- and ‘54149-CaSR complexes

To understand the molecular basis of recognition and to template subsequent optimization, we determined structures of CaSR in complex with two PAMs, **‘6218** and **‘54159** (*R-***‘85339**), derived from the 1.2 billion and the 2.7 million molecule screens, respectively. For CaSR-**‘6218** complex, the map was determined at a global nominal resolution of 2.8 Å with locally refined maps at resolutions of 2.7 Å and 3.4 Å for ECD-linker and linker–7TM regions, respectively (**fig. S4 and fig. S5)**. For CaSR- **‘54149** complex, the map was determined at a global nominal resolution of 2.7 Å with locally refined maps at resolutions of 2.6 Å and 3.6 Å for ECD-linker and linker–7TM regions, respectively (**fig. S4 and fig. S5**). Similar to the structures of cinacalcet- and evocalcet-bound CaSR complexes^16^, the 7TMs between two protomers adopt an asymmetric arrangement characterized by a raised position adopted by the TM6 of 7TM^A^ relative to the opposing TM6 of 7TM^B^ (**fig. S6. A to B**).

In the CaSR-**‘6218** complex, the PAM binding sites show density of **‘6218** in “extended” and “bent” conformations, recapitulating those of cinacalcet and evocalcet (**Fig. 3A, 3C**) (*22*). **‘6218** interacts with the same overall residues in both monomers, making conserved as well as site-specific interactions. In both sites, the cationic amine of the PAM ion-pairs with E837^7.32^ and hydrogen-bonds with Q681^3.33^. In the 7TM^B^ site, the methyl-benzazepine ring forms pi-pi interactions with F684^3.36^ and W818^6.50^, recapitulating the interactions formed by the naphthalene in cinacalcet and evocalcet. The benzoisothiazole ring makes pi-pi interactions with F821^6.53^ and Y825^6.57^ (**Fig. 3C**). In the 7TM^A^ site, while W818^6.50^ swings out by 120° and Y825^6.57^ moves down, the pi-pi interactions are still maintained. Conversely, the interaction with F821^6.53^ is lost as it swings out and is no longer part of the allosteric pocket (**Fig. 3A**). The docking predicted pose for **‘6218** superposes well with its experimental structure in both monomers (**Fig. 3B, 3D**). Both the docked and experimental poses of **‘6218** adopt an “extended” conformation in the 7TM^A^ site (**Fig. 3B**), while they have a “bent” conformation in the 7TM^B^ site (**Fig. 3D**). The same bent and extended conformations were observed for the initial docking hits; in this sense, this level of geometric fidelity emerged directly from the docking screen (**Fig. 1E, fig. S1**). The docked and experimental structures superimposed with a 1.88 Å root mean square deviation (RMSD) in the bent conformation monomer, and with a 2.23 Å RMSD in the extended conformation monomer. While the experimental results broadly support the docking prediction, there were important differences in the receptor structures. Compared to the cinacalcet complex against which we docked (7TM^B^), F821^6.53^ swings 120° into the site to become part of the binding pocket, making pi-pi interaction with the benzoisothiazole ring of **‘6218 (Fig. 3D**). This conformation is not adopted in the cinacalcet or the evocalcet complex, likely because the mobile groups of cinacalcet (1-propyl-3-(trifluoromethyl) benzene) and evocalcet (2-phenylacetic acid) are bulkier and would clash with this phenylalanine (**fig. S7**). Meanwhile, in the “extended” monomer’s binding site (7TM^A^), W818^6.50^ moves 120° to swing outside of the binding pocket in the **‘6218** complex.

Similar to **‘6218**, **‘54149** also adopts an “extended” conformation in the 7TM^A^ site and a “bent” conformation in the 7TM^B^ site, inducing similar rearrangements of W818^6.50^ and F821^6.53^ in the 7TM^A^ and 7TM^B^ site, respectively (**Fig. 3, E to H**). **‘54149** and **‘6218** share a benzoisothiazole group that is flexible in the two sites, suggesting the conformational changes of W818^6.50^ and F821^6.53^ are benzoisothiazole specific. At the 7TM^B^ site, the benzodioxole group interacts with F684^3.36^ and W818^6.50^ through pi-pi stacking, while the benzoisothiazole forms pi-pi interactions with F821^6.53^ and Y825^6.57^. The cationic amine hydrogen bonds with Q681^3.33^ and ion-pairs with E837^7.32^, and the adjacent methyl packs with I841^7.36^ (**Fig. 3G**). The interactions with F821^6.53^ and Y825^6.57^ are lost in the 7TM^A^ site as F821^6.53^ swings out of the pocket and Y825 swings down (**Fig. 3E**). The docked and experimental structures superposed to 0.91 Å RMSD in the 7TM^A^ site, and to 2.68 Å RMSD in the 7TM^B^ site (**Fig. 3, G to H**). Docking predicted **‘54149** to adopt both “extended” conformations in the binding pocket but we observe signs of conformational heterogeneity in the 7TM^A^ site. The EM density suggests that **‘54149** adopts an alternative “folded-over” conformation at this site, which has never been previously observed (**fig. S5C and fig. S8**). In this “folded-over” configuration, **‘54149** establishes favorable interactions with CaSR— the benzoisothiazole ring makes additional contacts by edge-to-pi stacking with F814^6.46^ and is surrounded by a hydrophobic pocket created by A840^7.35^, I841^7.36^, A844^7.39^ and V817^6.49^. Among those residues, A840^7.35^ and I841^7.36^ are important for the affinity of CaSR PAMs (*41, 42*). Unlike methyl-benzazepine (in **‘6218**) and naphthalene (in cinacalcet and evocalcet), **‘54149** employs a smaller benzodioxole as the stationary binding component, possibly allowing more configurations in the pocket. Together, the conformational disparity in the structure of these complexes highlights the ongoing importance of cycles of docking and structure determination in drug discovery efforts.

### ‘54149 stabilizes a distinct active-state CaSR dimer conformation

Compared to CaSR-cinacalcet alone, the structure against which we docked, our recent structure of the receptor in complex with cinacalcet and Gi*βγ* (PDB: 8SZH) (*43*) revealed that G protein binding promotes an additional conformational change that brings the two 7TMs into closer contact, in a configuration that is in line with the activation of other Family C receptors (*23, 44*). From the inactive state to the G-protein-bound active state, the interface contact area increases from 178.9 Å^2^ (inactive; NPS-2143 bound; PDB: 7M3E) to 206.2 Å^2^ (cinacalcet-bound; PDB: 7M3F) to 682.7 Å^2^ (cinacalcet, Gi*βγ* -bound) (calculated by PDBePISA). By aligning the 7TM^B^s, **‘54149** and **‘6218**’s 7TM^A^ moves down towards the cytoplasm associated with an increase in interface contact area to 351.3 and 271.5 Å compared to cinacalcet-bound CaSR, as illustrated by the relative positioning of the TM6 helices (**Fig. 4A**). The downward shift brings the two 7TMs in a conformation that is closer to the G protein-bound structure, especially for that induced by **‘54149**, suggesting that **‘54149** promotes a dimer configuration that may favor G-protein activation compared to those stabilized by the other compounds **(fig. S9)**. This may contribute to its efficacy and also potentially confer a different pharmacology.

**Fig. 4.**
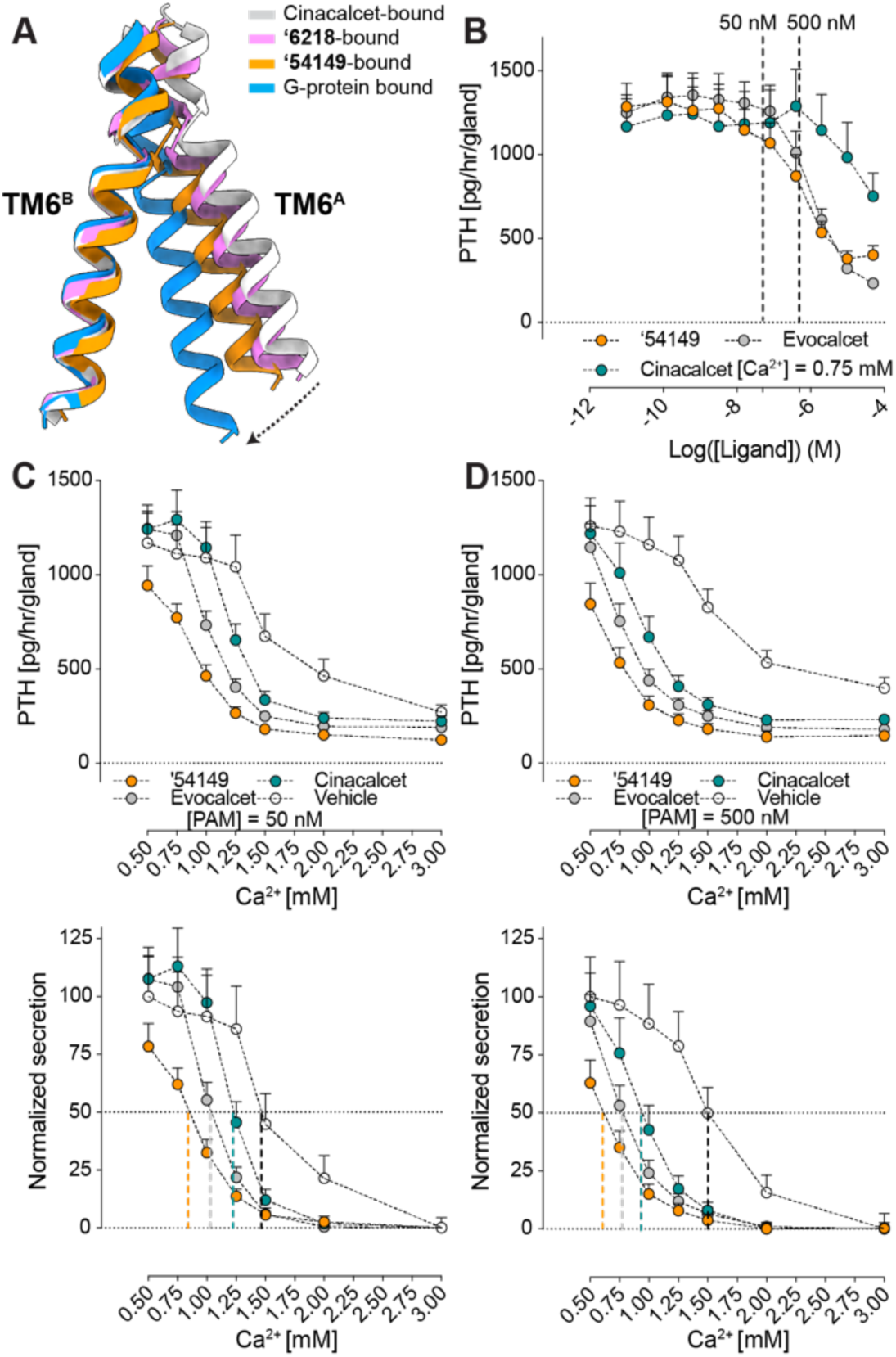
‘54149 increases the TM6-TM6 interface and is more effective in suppressing PTH secretion in *ex vivo* parathyroid glands. (**A**) The 7TM^A^ protomer undergoes a downward and rotational movement bringing TM6 closer to the 7TM^B^ from cinacalcet-bound to **‘54149**-bound structure to Gi-bound CaSR. Cinacalcet-bound CaSR is in grey, **‘54149**-bound CaSR is in orange, **‘6218**-bound CaSR is in pink and Gi-bound CaSR is in blue. (**B**) Parathyroid glands of 4-week-old C57/B6 wild-type (B6:Wt) mice were sequentially incubated with increasing concentrations of **‘54149**, cinacalcet and evocalcet from 0.01 nM to 50 *μ*M in the presence of 0.75 mM [Ca^2+^]_e_. The IC_50_s of **‘54149**, evocalcet and cinacalcet in suppressing PTH secretion are 583 nM [122 -4727 nM], 998 nM [412 – 4018 nM] and 53 μM respectively. (**C-D**) Parathyroid glands were sequentially incubated with increasing [Ca^2+^]_e_ from 0.5 mM to 3.0 mM in the presence of vehicle (0.1% DMSO), 50 nM (**C**) or 500 nM (**D**) of **‘54149**, cinacalcet or evocalcet. Top panels show changes in the rate of PTH secretion on a per-gland and per-hour basis with raising [Ca^2+^]_e_ to compare the PTH-max. Bottom panels show normalized PTH secretion rate (the highest rates are normalized to the basal rate at 0.5 mM [Ca^2+^]_e_ of the vehicle and the lowest rates are normalized to the rate at 3.0 mM [Ca^2+^]_e_) to better assess changes in the Ca^2+^-set-point ([Ca^2+^]_e_ needed to suppress 50% of [Ca^2+^]_e_-suppressible PTH secretion). Dotted vertical lines indicate Ca^2+^-set-points for the corresponding treatments. Mean ± SEM of n = 8 groups of PTGs for each treatment.

### ‘54149 suppresses PTH secretion better than the approved PAM drugs

Upon its activation, CaSR suppresses PTH secretion from parathyroid glands (*45*), which is the primary target of calcimimetic drugs. Since all PAM-bound structures were obtained under saturating calcium concentrations (10 mM), the different conformations observed are specific to each PAM and may be reflected in measurable functional differences. We thus investigated the functional effects of the different PAM drugs and leads by monitoring PTH secretion in extracted parathyroid glands from wild-type (WT) C57/BL6 (B6) mice at a constant external calcium concentration of 0.75 mM. All three of ‘**54149**, cinacalcet, and evocalcet inhibit PTH secretion dose-dependently, with potencies of **‘54149** (583 nM) ∼ evocalcet (998 nM) >> cinacalcet (53 μM) (**Fig. 4B**). As PAMs positively regulate CaSR by lowering the required calcium for activation, we wanted to assess how the different compounds shift the calcium set point for PTH secretion by the glands (**Fig. 4, C to D**). For this assay we used two PAM concentrations, 500 and 50 nM, (dashed line in **Fig. 4B**). At 500 nM, **‘54149** shifted the calcium set point from 1.5 mM to 0.62 mM, while at the same concentration, cinacalcet shifted the set point to ∼0.94 mM and evocalcet shifted it to 0.76 mM (**Fig. 4D**). The same trend holds when the PAMs were administered at 50 nM, leading to shifts in the calcium set-point from 1.47 (vehicle) to 1.23 (cinacalcet) to 1 (evocalcet) to 0.85 (**’54149**) mM (**Fig. 4C**). It is worth noting that **‘54149** also suppresses the tonic secretion of PTH at 0.5 mM calcium, an effect not observed with the two approved drugs.

### ‘54149 reduces serum PTH at lower doses with less hypocalcemia than cinacalcet

Encouraged by its improved affinity and *ex vivo* organ efficacy, we investigated the *in vivo* activity of ‘**54149**, beginning with pharmacokinetic (PK) studies in CD-1 mice. We administered ‘**54149** at a dose of 3 mg/kg subcutaneously, and dosed cinacalcet and evocalcet in the same manner for direct comparison (**Fig. 5A**). At this dose, ‘**54149** was found in appreciable amount in plasma— AUC_0→inf_ 18,500 mg*min/ml. The C_max_ reaches 112 ng/ml (**340 nM**) at 15 min and stays high until 60 min (100 ng/ml). Based on ‘**54149**’s EC_50_, ‘**54149** is close to saturation at 3 mg/kg dose over this period (**Fig. 2E**). By comparison, evocalcet has a much higher systemic exposure at the same dose, with a C_max_ of 3,250 ng/ml (**8.68 μM**) at 60 min. On the other hand, cinacalcet - which is far more widely used - has lower exposure than ‘**54149**, with C_max_ of 58.9 ng/ml (**149.5 nM**) 15 min after subcutaneous administration (**Fig. 5A**). We note that no effort has been made to optimize ‘**54149** for pharmacokinetic exposure or clearance—to the extent that it has favorable PK, this simply reflects the physical property constraints imposed in docking and ligand optimization.

**Fig. 5.**
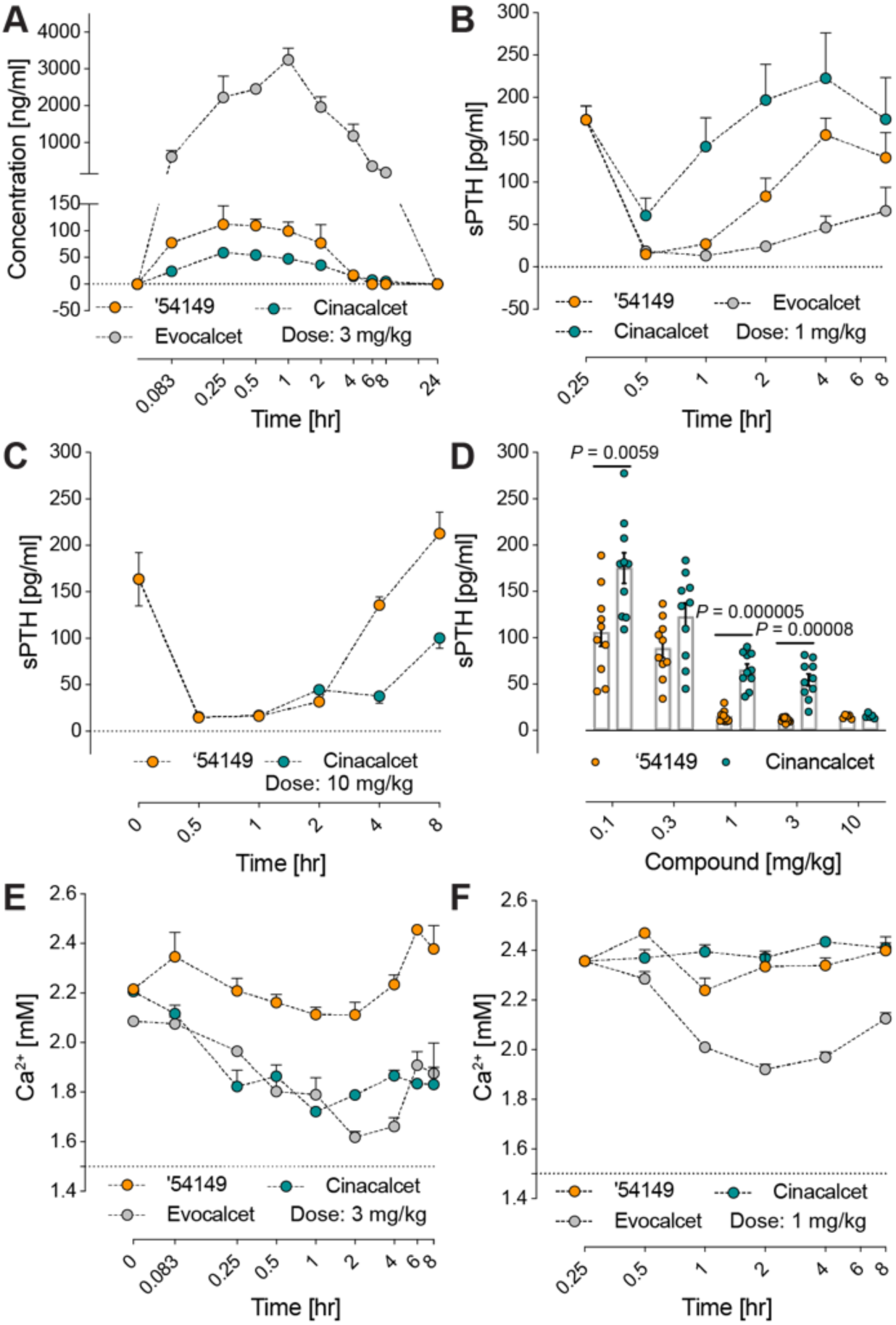
‘54149 suppresses serum PTH at lower dose and causes less hypocalcemia effect than cinacalcet and evocalcet. (**A**) Pharmacokinetics of **‘54149** compared to cinacalcet and evocalcet after 3 mg/kg subcutaneous injection. (**B**) Serum PTH concentration change over 8 hours after 1 mg/kg subcutaneous injection of **‘54149**, cinacalcet or evocalcet. (**C**) Serum PTH concentration change over 8 hours after 10 mg/kg subcutaneous injection **‘54149** or cinacalcet (n = 5). (**D**) Comparison of **‘54149** to cinacalcet in regulating serum PTH at different doses (subcutaneous injection) after 30min of injection. Each dose consists of n = 10 mice except injection at 10 mg/kg (n = 5). P-values were assessed by unpaired Student’s t-test. (**E**) Plasma calcium concentration in mice after 3 mg/kg subcutaneous injection of **‘54149**, cinacalcet or evocalcet. (**F**) Serum calcium concentration after 1 mg/kg subcutaneous injection of **‘54149**, cinacalcet or evocalcet. For experiments in panel **B-D**, **F**, the concentrations of evocalcet and cinacalcet are corrected for their molecular weight difference with **‘54149**.

Based on the PK, we picked two doses to investigate the time course of PTH suppression by the PAMs in WT B6 mice. At 1 mg (3.1 µmol/kg), ‘**54149** and equimolar evocalcet fully suppress PTH secretion, while cinacalcet is less effective at this dose (**Fig. 5B**). Only at 10 mg/kg (31 µmole/kg) was cinacalcet able to fully suppress PTH secretion (**Fig. 5, C to D**). Overall, **‘54149** fully suppresses serum PTH at 10 times lower dose than cinacalcet (**Fig. 5D**), consistent with its ability to suppress releases of both tonic and Ca^2+^-suppressible pools of PTH (**Fig. 4, C to D**).

A key adverse effect of cinacalcet and etelcalcetide is decreased blood calcium (*46*). In secondary hyperparathyroidism (SHPT), high PTH is accompanied by low or normal blood calcium concentration. The overproduction of PTH and the proliferation of parathyroid cells in patients with SHPT are largely driven by low blood calcium and high blood phosphate levels (*47–49*) as well as reduced CaSR expression in parathyroid cells (*50*). We were thus keen to compare the serum calcium concentration after injection of ‘**54149** versus cinacalcet. At the dose of 3mg/kg, ‘**54149** did not significantly alter serum calcium concentration for 4 hrs, but slightly increased it from 2.2 to 2.4 mM after the drug dissipated in circulation 6 hrs post-injection (**Fig. 5E**). In contrast, the same dose of cinacalcet and evocalcet significantly lowered serum calcium for more than 8 hrs from 2.2 mM to the lowest levels of 1.7 mM and 1.6 mM, respectively. The hypocalcemic action of evocalcet is particularly robust even at a lower dose of 3.1 µmol/kg (∼1 mg/kg) (**Fig. 5F**), while the same dose of **‘54149** retained the ability to maximally suppress serum PTH without producing hypocalcemia (**Fig. 5D**). Although the mechanisms underlying the different calcemic actions of these 3 compounds remain to be determined, their common ability to suppress PTH secretion suggests that differential calcemic actions likely take place in other calciotropic organs outside of parathyroid glands.

## Discussion

Four key observations emerge from this study. **First**, from a structure-based screen of a 1.2 billion molecule tangible library emerged a spectrum of diverse chemotypes that potently enhanced CaSR activation. The new molecules represent among the first positive allosteric modulators (PAMs) discovered via large library docking, and among the first structure-based ligands discovered for Family C GPCRs. The potency of the initial docking hits was relatively high, with EC_50_ values down to 270 nM, and all were topologically dissimilar to known CaSR PAMs. Structure-based optimization improved affinity between 40 and 600-fold, leading to molecules that were up to 50-fold more potent than cinacalcet *in vitro* and 10 to 100-fold more potent at suppressing PTH secretion from organs *ex vivo* as well as *in vivo* in animals. **Second**, the docking predictions were largely confirmed by the subsequent cryo-EM structures, with an important exception (see below), including selecting for and correctly predicting extended and bent conformations in the TM^A^ and TM^B^ sites of the CaSR dimer. **Third,** our direct comparison for the impact of docking an ultra-large (1.2 billion) library versus a smaller (2.7 million) molecule library in the same pocket shows the improvement in docking scores as the library size increases, an effect that has been previously suggested by simulations (*40*) but not experimentally tested in a controlled way (**Fig. 1C**), Here, experimental docking hit rates were 2.7-fold higher in the large library screen than in the “in-stock” screen, and the best hits from the large library were up to 37-fold more potent. **Fourth,** the new chemotypes make new interactions with the receptor, promoting new active-state dimer interfaces that are closer to the G protein coupled state which were not observed with the established drugs. In this sense, the experimental structures provide an additional layer of information in terms of global conformations that may help explain differences in the relative efficacy and pharmacology of different ligands. Correspondingly, **‘54149** promotes a TM6-TM6 interface that is closest to the fully active G protein-coupled state of the receptor dimer and is highly potent in suppressing PTH secretion, while also seemingly devoid of the hypocalcemia that is the key dose-limiting side effect of approved calcimimetic drugs (*51, 52*).

Several caveats merit mentioning. We do not pretend the molecules described here are drugs. Whereas the pharmacokinetics of **‘54149** are sufficient to support *in vivo* studies, and indeed in some ways to demonstrate superiority to cinacalcet, there is clearly room for optimization of exposure and half-life of the molecule. While the relative lack of a hypocalcemic effect is encouraging, understanding the mechanism underlying this effect requires systematic exploration of CaSR activation in other calciotropic organs, including bones and kidneys. Further, whereas in three of the four cases the docking predicted structures of the PAMs in the 7TM^A^ and 7TM^B^ monomers were confirmed by cryo-EM, in one site the docking pose was different from the experimental result. Finally, while the improvement in docking hit rates and docking potencies from billion molecules versus million molecule libraries seems compelling, the numbers experimentally tested remain relatively low given docking uncertainties.

These caveats should not obscure the main observations: From docking 1.2 billion molecules against the structure of CaSR emerged potent new positive allosteric modulators, topologically dissimilar to the known ligands of this receptor. Structure-based optimization of the new PAMs led to molecules with *in vitro* potencies in the low nM range, up to 14-fold more potent than the standard of care for the calcimimetic drugs, cinacalcet. In *ex-vivo* organ studies this increase in potency was retained, while *in vivo*, too, the new molecules were also substantially more potent than cinacalcet. The novel chemotypes stabilized CaSR dimer conformations that are not observed in the previous structures of established PAMs, which may underlie the ability of the new chemotypes to support strong efficacy in suppressing parathyroid hormone secretion without inducing their dose-limiting hypocalcemia. Finally, docking hits were 37-fold more potent, and docking hit-rates 2.7-fold higher in the billion-molecule library campaign than for docking the million-molecule scale library against the same site. While such a comparison merits further study, certainly with more molecules being tested, it is consistent with theoretical studies (*40*) and supports the continued expansion of readily testable libraries for drug discovery (*26, 29*).

## Funding

National Institutes of Health grant R01NS122394 (GS)

National Institutes of Health grant R01DK132902 (GS)

National Institutes of Health grant R35GM122481 (BKS)

National Institutes of Health grant R35GM71896 (JJI)

National Institutes of Health grant RF1AG075742 (WC)

National Institutes of Health grant R01DK122259 (WC)

BLR&D I01BX005851 (WC)

BLR&D IK6BX004835 (WC)

Damon Runyon Postdoctoral Research Fellowship (FL)

## Author Contributions

FL conducted the docking screens and the ligand optimization with assistance from ALK and JL, advised by BKS. CGW conducted the *in vitro* activity assays, with early assistance from JM, and determined the structures by cryo-EM, advised by GS. CLT, ZC, and WC conducted *ex vivo* and *in vivo* activity assays. Aggregation studies were conducted by IG. JJI developed and prepared the make-on-demand library assisted with large library docking strategies. OOT and YSM supervised compound synthesis of Enamine compounds purchased from the ZINC22 database and the 46 billion catalog library.

## Data and materials availability

DOCK3.7 and DOCK3.8 are available without charge for academic use https://dock.compbio.ucsf.edu/. Most underlying data from this study are included among the primary figures and tables, and in the SI, any not so included are available from the authors on request. All molecules tested are available from Enamine and may be accessed via their ZINC numbers (SI Tables 1). Plasmids and reagents to conduct BRET signaling assays are available from GS. Mouse lines are available from Jackson Laboratory. Cryo-EM structures and maps are available in the Protein Data Bank and EMDB under accession numbers PDBID XXXX, PDBID FFFF, and EMDB YYYY, EMDB GGGG, respectively.

## References

1. E. M. Brown et al., Cloning and characterization of an extracellular Ca(2+)-sensing receptor from bovine parathyroid. Nature 366, 575–580 (1993).

2. K. Leach et al., International Union of Basic and Clinical Pharmacology. CVIII. Calcium-Sensing Receptor Nomenclature, Pharmacology, and Function. Pharmacol Rev 72, 558–604 (2020).

3. F. M. Hannan, E. Kallay, W. Chang, M. L. Brandi, R. V. Thakker, The calcium-sensing receptor in physiology and in calcitropic and noncalcitropic diseases. Nat Rev Endocrinol 15, 33–51 (2018).

4. F. M. Hannan, R. V. Thakker, Calcium-sensing receptor (CaSR) mutations and disorders of calcium, electrolyte and water metabolism. Best Pract Res Clin Endocrinol Metab 27, 359–371 (2013).

5. M. R. Pollak et al., Autosomal dominant hypocalcaemia caused by a Ca(2+)-sensing receptor gene mutation. Nat Genet 8, 303–307 (1994).

6. F. M. Hannan et al., Identification of 70 calcium-sensing receptor mutations in hyper- and hypo-calcaemic patients: evidence for clustering of extracellular domain mutations at calcium-binding sites. Hum Mol Genet 21, 2768–2778 (2012).

7. S. H. Pearce et al., A familial syndrome of hypocalcemia with hypercalciuria due to mutations in the calcium-sensing receptor. N Engl J Med 335, 1115–1122 (1996).

8. J. Patel, M. B. Bridgeman, Etelcalcetide (Parsabiv) for Secondary Hyperparathyroidism in Adults With Chronic Kidney Disease on Hemodialysis. P T 43, 396–399 (2018).

9. L. Pereira, C. Meng, D. Marques, J. M. Frazao, Old and new calcimimetics for treatment of secondary hyperparathyroidism: impact on biochemical and relevant clinical outcomes. Clin Kidney J 11, 80–88 (2018).

10. T. C. Sauter et al., Calcium Disorders in the Emergency Department: Independent Risk Factors for Mortality. PLoS One 10, e0132788 (2015).

11. Z. Zhang, X. Xu, H. Ni, H. Deng, Predictive value of ionized calcium in critically ill patients: an analysis of a large clinical database MIMIC II. PLoS One 9, e95204 (2014).

12. M. Egi et al., Ionized calcium concentration and outcome in critical illness. Crit Care Med 39, 314–321 (2011).

13. T. Steele, R. Kolamunnage-Dona, C. Downey, C. H. Toh, I. Welters, Assessment and clinical course of hypocalcemia in critical illness. Crit Care 17, R106 (2013).

14. A. Husain, R. J. Simpson, Jr., G. Joodi, Serum Calcium and Risk of Sudden Cardiac Arrest in the General Population. Mayo Clin Proc 93, 392 (2018).

15. R. Nardone, F. Brigo, E. Trinka, Acute Symptomatic Seizures Caused by Electrolyte Disturbances. J Clin Neurol 12, 21–33 (2016).

16. T. B. Drueke, Cell biology of parathyroid gland hyperplasia in chronic renal failure. J Am Soc Nephrol 11, 1141–1152 (2000).

17. J. C. Bureo et al., Prevalence of secondary hyperparathyroidism in patients with stage 3 and 4 chronic kidney disease seen in internal medicine. Endocrinol Nutr 62, 300–305 (2015).

18. A. Levin et al., Prevalence of abnormal serum vitamin D, PTH, calcium, and phosphorus in patients with chronic kidney disease: results of the study to evaluate early kidney disease. Kidney Int 71, 31–38 (2007).

19. D. L. Andress et al., Management of secondary hyperparathyroidism in stages 3 and 4 chronic kidney disease. Endocr Pract 14, 18–27 (2008).

20. C. P. Kovesdy, Epidemiology of chronic kidney disease: an update 2022. Kidney Int Suppl (2011) 12, 7–11 (2022).

21. J. P. Pin, T. Galvez, L. Prezeau, Evolution, structure, and activation mechanism of family 3/C G-protein-coupled receptors. Pharmacol Ther 98, 325–354 (2003).

22. Y. Gao et al., Asymmetric activation of the calcium-sensing receptor homodimer. Nature 595, 455–459 (2021).

23. A. B. Seven et al., G-protein activation by a metabotropic glutamate receptor. Nature 595, 450–454 (2021).

24. M. M. Papasergi-Scott et al., Structures of metabotropic GABA(B) receptor. Nature 584, 310–314 (2020).

25. J. Lyu et al., Ultra-large library docking for discovering new chemotypes. Nature 566, 224–229 (2019).

26. C. Gorgulla et al., An open-source drug discovery platform enables ultra-large virtual screens. Nature 580, 663–668 (2020).

27. R. M. Stein et al., Virtual discovery of melatonin receptor ligands to modulate circadian rhythms. Nature 579, 609–614 (2020).

28. A. Alon et al., Structures of the sigma(2) receptor enable docking for bioactive ligand discovery. Nature 600, 759–764 (2021).

29. A. A. Sadybekov et al., Synthon-based ligand discovery in virtual libraries of over 11 billion compounds. Nature 601, 452–459 (2022).

30. E. A. Fink et al., Structure-based discovery of nonopioid analgesics acting through the alpha(2A)-adrenergic receptor. Science 377, eabn7065 (2022).

31. I. Singh et al., Structure-based discovery of conformationally selective inhibitors of the serotonin transporter. Cell 186, 2160–2175 e2117 (2023).

32. R. G. Coleman, M. Carchia, T. Sterling, J. J. Irwin, B. K. Shoichet, Ligand pose and orientational sampling in molecular docking. PLoS One 8, e75992 (2013).

33. E. C. Meng, B. K. Shoichet, I. D. Kuntz, Automated Docking with Grid-Based Energy Evaluation. J Comput Chem 13, 505–524 (1992).

34. K. A. Sharp, R. A. Friedman, V. Misra, J. Hecht, B. Honig, Salt effects on polyelectrolyte-ligand binding: comparison of Poisson-Boltzmann, and limiting law/counterion binding models. Biopolymers 36, 245–262 (1995).

35. K. Gallagher, K. Sharp, Electrostatic contributions to heat capacity changes of DNA-ligand binding. Biophys J 75, 769–776 (1998).

36. M. M. Mysinger, B. K. Shoichet, Rapid context-dependent ligand desolvation in molecular docking. J Chem Inf Model 50, 1561–1573 (2010).

37. S. Gu, M. S. Smith, Y. Yang, J. J. Irwin, B. K. Shoichet, Ligand Strain Energy in Large Library Docking. J Chem Inf Model 61, 4331–4341 (2021).

38. R. H. J. Olsen et al., TRUPATH, an open-source biosensor platform for interrogating the GPCR transducerome. Nat Chem Biol 16, 841–849 (2020).

39. B. I. Tingle et al., ZINC-22 horizontal line A Free Multi-Billion-Scale Database of Tangible Compounds for Ligand Discovery. J Chem Inf Model 63, 1166–1176 (2023).

40. J. Lyu, J. J. Irwin, B. K. Shoichet, Modeling the expansion of virtual screening libraries. Nat Chem Biol 19, 712–718 (2023).

41. K. Leach et al., Towards a structural understanding of allosteric drugs at the human calcium-sensing receptor. Cell Res 26, 574–592 (2016).

42. A. N. Keller et al., Identification of Global and Ligand-Specific Calcium Sensing Receptor Activation Mechanisms. Mol Pharmacol 93, 619–630 (2018).

43. F. He et al., Allosteric modulation and G-protein selectivity of the Ca^2+^-sensing receptor. Nature, (2024). Feb 7. doi: 10.1038/s41586-024-07055-2. Epub ahead of print. PMID: 38326620.

44. S. Lin et al., Structures of G(i)-bound metabotropic glutamate receptors mGlu2 and mGlu4. Nature 594, 583–588 (2021).

45. E. M. Brown, Clinical lessons from the calcium-sensing receptor. Nat Clin Pract Endocrinol Metab 3, 122–133 (2007).

46. G. A. Block et al., Effect of Etelcalcetide vs Cinacalcet on Serum Parathyroid Hormone in Patients Receiving Hemodialysis With Secondary Hyperparathyroidism: A Randomized Clinical Trial. JAMA 317, 156–164 (2017).

47. S. A. Jamal, P. D. Miller, Secondary and tertiary hyperparathyroidism. J Clin Densitom 16, 64–68 (2013).

48. M. Rodriguez, E. Nemeth, D. Martin, The calcium-sensing receptor: a key factor in the pathogenesis of secondary hyperparathyroidism. Am J Physiol Renal Physiol 288, F253–264 (2005).

49. P. P. Centeno et al., Phosphate acts directly on the calcium-sensing receptor to stimulate parathyroid hormone secretion. Nat Commun 10, 4693 (2019).

50. J. Gogusev et al., Depressed expression of calcium receptor in parathyroid gland tissue of patients with hyperparathyroidism. Kidney Int 51, 328–336 (1997).

51. G. S. Schmidt, T. D. Weaver, T. D. Hoang, M. K. M. Shakir, Severe Symptomatic Hypocalcemia, complicating cardiac arrhythmia following Cinacalcet (Sensipar(TM)) administration: A Case Report. Clin Case Rep 9, e04876 (2021).

52. G. A. Block et al., Cinacalcet for secondary hyperparathyroidism in patients receiving hemodialysis. N Engl J Med 350, 1516–1525 (2004).

53. J. M. Word, S. C. Lovell, J. S. Richardson, D. C. Richardson, Asparagine and glutamine: using hydrogen atom contacts in the choice of side-chain amide orientation. J Mol Biol 285, 1735–1747 (1999).

54. G. M. Sastry, M. Adzhigirey, T. Day, R. Annabhimoju, W. Sherman, Protein and ligand preparation: parameters, protocols, and influence on virtual screening enrichments. J Comput Aided Mol Des 27, 221–234 (2013).

55. K. A. Sharp, Polyelectrolyte Electrostatics - Salt Dependence, Entropic, and Enthalpic Contributions to Free-Energy in the Nonlinear Poisson-Boltzmann Model. Biopolymers 36, 227–243 (1995).

56. R. M. Stein et al., Property-Unmatched Decoys in Docking Benchmarks. J Chem Inf Model 61, 699–714 (2021).

57. A. V. Fassio et al., Prioritizing Virtual Screening with Interpretable Interaction Fingerprints. J Chem Inf Model 62, 4300–4318 (2022).

58. D. N. Mastronarde, Automated electron microscope tomography using robust prediction of specimen movements. J Struct Biol 152, 36–51 (2005).

59. A. Punjani, J. L. Rubinstein, D. J. Fleet, M. A. Brubaker, cryoSPARC: algorithms for rapid unsupervised cryo-EM structure determination. Nat Methods 14, 290–296 (2017).

60. J. Zivanov et al., New tools for automated high-resolution cryo-EM structure determination in RELION-3. Elife 7, (2018).

61. E. F. Pettersen et al., UCSF Chimera--a visualization system for exploratory research and analysis. J Comput Chem 25, 1605–1612 (2004).

62. P. Emsley, B. Lohkamp, W. G. Scott, K. Cowtan, Features and development of Coot. Acta Crystallogr D Biol Crystallogr 66, 486–501 (2010).

63. D. Liebschner et al., Macromolecular structure determination using X-rays, neutrons and electrons: recent developments in Phenix. Acta Crystallogr D Struct Biol 75, 861–877 (2019).

64. V. B. Chen et al., MolProbity: all-atom structure validation for macromolecular crystallography. Acta Crystallogr D Biol Crystallogr 66, 12–21 (2010).

65. E. F. Pettersen et al., UCSF ChimeraX: Structure visualization for researchers, educators, and developers. Protein Sci 30, 70–82 (2021).

66. W. Chang, C. Tu, T. H. Chen, D. Bikle, D. Shoback, The extracellular calcium-sensing receptor (CaSR) is a critical modulator of skeletal development. Sci Signal 1, ra1 (2008).

67. W. Chang et al., PTH hypersecretion triggered by a GABA(B1) and Ca(2+)-sensing receptor heterocomplex in hyperparathyroidism. Nat Metab 2, 243–255 (2020).

