## Supplementary Materials for "Small vs. Large Library Docking for Positive Allosteric Modulators of the Calcium Sensing Receptor"

**The PDF file includes:**

Materials and Methods

Figs. S1 to S8

Tables S1 to S2

References

`

#### Materials and Methods

##### In-stock and ultra-large virtual ligand screening

To investigate the effect of small versus large library docking and test the docking prediction of the positive allosteric modulator (PAM) binding sites in complex with "extended" or "bent" PAMs, we optimized two docking set ups based on the cryo-EM structures of cinacalcet- or evocalcet-bound CaSR. CaSR/cinacalcet (PDB: 7M3F) is used for $7TM^A$ site, and CaSR/evocalcet (PDB: 7M3G) is used for $7TM^B$ site (*22*). In both sites, the position of Q681 and E837 are manually adjusted to form stronger hydrogen bonds or salt bridge with the secondary amine in cinacalcet or evocalcet, and in the $7TM^B$ site, lipid tails were added in the docking set up based on the existing electron density. 7TMs were protonated using Reduce (*53*) ($7TM^B$ site) or by Protein Preparation Wizard in Maestro ($7TM^A$ site) (2020 release) (*54*). Energy grids for the different energy terms of the scoring function were pre-generated--van der Waals term based on the AMBER force fields using CHEMGRID (*33*); Poisson–Boltzmann-based electrostatic potentials using QNIFFT73 (*35, 55*); context-dependent ligand desolvation was calculated using SOLVMAP (*36*). The volume of the low dielectric and the desolvation volume was extended out 0.8 and 0.3 Å in $7TM^A$ site and 0.6 and 0.3 Å in $7TM^B$ site. The experimentally determined poses of cinacalcet and evocalcet were used to generate matching spheres, which are later used by the docking software to fit pre-generated ligands' conformations into the small molecule binding sites (*32*).

The resulting docking set-ups were evaluated for its ability to enrich known CaSR ligands over property-matched decoys. Decoys are theoretical non-binders to the receptor as they are topologically dissimilar to known ligands but retain similar physical properties. We extracted 10 known PAMs from CHEMBL (https://www.ebi.ac.uk/chembl/) including cinacalcet and evocalcet. Four-hundred and eighty-five decoys were generated by using the DUDE-Z pipeline (*56*). high logAUCs of 38.89 and 31.67 were achieved for $7TM^A$ site and $7TM^B$ site respectively. Moreover, these docking set-ups offer fidelity in reproducing "extended" and "bent" poses of the known PAMs. For example, by using the $7TM^A$ site set-up, 7 out of 10 PAMs adopt an "extended" conformations, while making sensible interactions with the surrounding key residues. By using the $7TM^{A}$ site set-up, 7 out of 10 PAMs adopt an "bent" conformations. We also used "extrema" set of 92,552 molecules using the DUDE-Z web server (http://tldr.docking.org) to ensure that the set ups do not enrich extreme physical properties. Both set ups enrich over 90% neutrals or mono-cations among the top-ranking molecules, which are two charges that have precedents of acting as CaSR PAMs.

2.7 million "lead-like" molecules (molecular weight 300-350 Da and logP $\leq$ 3.5), from ZINC15 database (http://zinc15.docking.org/), were docked against both sites using DOCK3.7 (*32*). In the docking screen against the $7TM^{A}$ site, each library molecule was sampled in about 3,927 orientations and, on average, 330 conformations. For the $7TM^{B}$ site, each library molecule was sampled in about 3,612 orientations and, on average, 330 conformations. The best scoring configuration for each docked molecule was relaxed by rigid-body minimization. The two screens took 956 and 917 core hours respectively spread over 100 cores, or slightly more than 3 days. For the 1.2-billion ultra-large library docking, each library from the ZINC22 database (*39*) was sampled in about 1,707 orientations and 425 conformations in the $7TM^{B}$ site by using DOCK3.8 (*32*). Overall, over 681 trillion complexes were sampled and scored, spending 380,016 core hours spreading over 2,000 cores, or around 7 days.

**Docking results' processing**

For the in-stock screen against the $7TM^{B}$ site, 5,208 molecules with dock energy $\leq$ -35 kcal/mol were filtered for novelty using the ECP4-based Tanimoto coefficient (Tc) against 662 CaSR ligands in CHEMBL (https://www.ebi.ac.uk/chembl/). Molecules with Tc > 0.35 were eliminated. These molecules are filtered for internal strains with criteria of total strain energy < 8 and maximum dihedral torsion energy < 3 (*37*). Moreover, the molecules are further filtered for key interactions: hydrogen bond with Q681, salt bridge with E837 by interfilter.py based on OpenEye

Python Toolkits (https://docs.eyesopen.com/toolkits/python/quickstart-python/linuxosx.html). After these three filters, 103 molecules were left for further examination. Upon clustering by an ECP4-based *Tc* of 0.5, 79 molecules were visually inspected for pi-pi interactions with W818 and F684. 28 molecules were picked, but only 22 molecules can be sourced from vendors and arrived for *in vitro* testing.

For the in-stock screen against the 7TM$^{A}$ site, 33,321 molecules with dock energy $\leq$ -43 kcal/mol were filtered against the same three filters, resulting in 2,540 molecules for further examination. The 2,540 molecules were filtered against a vendor filter to assess their persuasibility, resulting in 647 molecules for further examination. The 647 molecules were clustered based on ECP4-based *Tc* of 0.5 and result in 413 clusterheads. The clusterheads were visually inspected in a similar manner resulting in 28 candidates ordering for purchasing, and 26 molecules arrived for testing. For the large-scale screen, 1.2 billion molecules were screened, and 1 billion molecules scored in the 7TM$^{B}$ site. The strain filter is incorporated as part of the new DOCK3.8 pipeline. 2,321,171 molecules with $\leq$ -35 kcal/mol were filtered for key interactions with Q681, E837, W818 and F684 and novelty. The interaction filtering script for pi-pi interactions with W818 and F684 is implemented based on LUNA (https://github.com/keiserlab/LUNA) (*57*). After visual inspection, 1,002 molecules were left. To reduce the number of candidate molecules for purchasing, these 1,002 molecules were re-docked against the 7TM$^{A}$ site, and 907 molecules were scored in the 7TM$^{A}$ site. To the end, the molecules were visually inspected again for their poses against both sites, and the remaining 212 novel and non-strained molecules were clustered by the LUNA 1,024-length binary fingerprint of a *Tc* = 0.3, resulting in 112 clusterheads. Ultimately, 96 molecules were prioritized for purchasing based on a final round of visual inspection. The 96 molecules belong to three categories—(1) molecules that have 2 aromatic ends, and they usually adopt "bent" pose in 7TM$^{B}$ site and "extended" pose in 7TM$^{A}$ site. (2) molecules that have aromatic moiety in the pocket and non-ring structure at the distal end but scores well. (3) interesting or neutral molecules.

#### Synthesis of molecules

The in-stock prioritized molecules were sourced from Enamine, Vitas-M laboratory, Ltd., UkrOrgSynthesis Ltd., ChemBridge Corporation and Sigma. Ninety-six molecules prioritized for purchasing were synthesized by Enamine for a total fulfilment rate of 74%. Compounds were sourced from the Enamine REAL database (https://enamine.net/compound-collections/real-compounds). The purities of active molecules were at least 90% and typically above 95%. The detailed chemical synthesis can be found in the Chemical Synthesis and analytical investigations section.

#### Hit Optimization

Potential analogs of the hits were identified through a combination of similarity and substructure searches of the SmallWorld (https://sw.docking.org/) from a 46 billion make-on-demand library. Potential analogs were docked to the CaSR 7TM$^{B}$ binding site using DOCK3.8. As was true in the primary screen, the resulting docked poses were manually evaluated for specific interactions and compatibility with the site, and prioritized analogs were acquired and tested experimentally.

#### Pharmacokinetics

Pharmacokinetic experiments of **'54149**, cinacalcet and evocalcet were performed by Bienta Enamine Biology Sciences (Kiev, Ukarine) in accordance with the Study Protocols P092622a, P050723b and P050723a. Plasma pharmacokinetics of **'54149**, cinacalcet and evocalcet were measured after a single 3 mg/kg dose, administered subcutaneously (SC) at time points of 5, 15, 30, 60, 120, 240, 360, 480 and 1,440 min. All animals were fasted for 4h before dosing. **'54149** was formulated in 2-HPbCD – saline (30%:70%, v/v). Cinacalcet and evocalcet were formulated

`

in DMSO – 20% Captisol in saline w/v (10:90, v/v). Testing was done in healthy male CD-1 mice (9 weeks old) weighing 32.7 ± 2.1 g, 32.8 ± 1.9 g or 32.9 ± 2.4 g in the three studies. For all three studies, each of the time point treatment group included 3 animals with a control group of one animal dosed with vehicle. In total, 28 animals were used in each study. Mice were injected IP with 2,2,2-tribromoethanol at the dose of 150 mg/kg prior to drawing the blood. Blood collection was performed from the orbital sinus in microtainers containing K3EDTA and tubes with clot activator. Animals were sacrificed by cervical dislocation after the blood samples collection. Blood samples were centrifuged for 10 min to obtain plasma (15 min to obtain serum) at 3000 rpm. All samples were immediately processed, flash-frozen and stored at -70°C until subsequent analysis. The concentrations of the test compound below the lower limits of quantitation (LLOQ = 2 ng/ml) were designated as zero. The pharmacokinetic data analysis was performed using noncompartmental, bolus injection or extravascular input analysis models in WinNonlin 5.2 (PharSight). Data below LLOQ were presented as missing to improve validity of T1/2 calculations. For each treatment condition, the final concentration values obtained at each time point were analyzed for outliers using Grubbs' test with the level of significance set at $p < 0.05$.

<u>Sample Processing:</u> Plasma samples (40 µl) were mixed with 200 µl of internal standard solution. After mixing by pipetting and centrifuging for 4 min at 6,000 rpm, 2 µl of each supernatant was injected into LC-MS/MS system. Solution of compound Verapamil (200 ng/ml in water-methanol mixture 1:9, v/v) was used as internal standard for quantification of **'54149** in plasma samples. Solution of Prometryn (100 ng/ml in water-methanol mixture 1:9, v/v) was used as internal standard for quantification of cinacalcet in plasma samples. Solution of Imipramine (50 ng/ml in water-methanol mixture 1:9, v/v) was used as internal standard for quantification of evocalcet in plasma samples.

`

Data Analysis: Peak plasma concentration ($C_{max}$) and time for the peak plasma concentration ($T_{max}$) were the observed values. The areas under the concentration time curve ($AUC_{last}$ and $AUC_{inf}$) were calculated by the linear trapezoidal rule. The terminal elimination rate constant, ke was determined by regression analysis of the linear terminal portion of the log plasma concentration-time curve. Mean, SD and %CV was calculated for each analyte.

Serum Calcium Measurement: Serum Calcium level was determined using commercial kits according to the manufacturer's instructions. The principle of the method is the ability of calcium forms a blue-colored complex with Arsenazo III dye at neutral pH, the intensity of which is proportional to the concentration of calcium. Interference with magnesium is eliminated by the addition of 8-hydroxyquinoline-5-sulfonic acid. Reproducibility: CV=2.91 %.

**Molecular cloning**

Full-length (FL) and the truncated CaSR (residues 20-894), were cloned into a pFastBac1 vector (for expression in insect cells) or a pcDNA3.1(+) vector (for expression in HEK293S cells), with a N-terminal haemagglutinin (HA) signal sequence followed by a FLAG tag. To improve the protein yield of CaSR, the DNA sequence of the C-terminal tail from $GABA_{B1}$ or $GABA_{B2}$ and an endoplasmic reticulum retention motif were inserted at the C-terminus of pFastBac1-FLAG-CaSR (20-894) to generate CaSR-C1 and CaSR-C2 constructs, which have been shown to have comparable G-protein signaling profiles as the WT CaSR homodimer[16]. The FLAG tag of CaSR-C1 construct was then replaced by a Twin-Strep-tag (WSHPQFEKGGGSGGGSGGSAWSHPQFEK). All plasmids used were sequence-verified.

**Bioluminescence Resonance Energy Transfer (BRET) TRUPATH Assay**

BRET assays were performed and analyzed similar to previously described methods (*38*). HEK-293S cells grown in FreeStyle 293 suspension media (ThermoFisher) were co-transfected with

`

150 ng of pCDNA3.1-CaSR FL, Gαi3-Rluc8, Gβ, and Gγ-GFP2 per 1ml of cells at a density of 1 x $10^6$ cells $ml^{-1}$ using a DNA/polyethylineimine ratio of 1:5, and incubated at 130 rpm., 37 °C. Cells were harvested 48 h post-transfection, washed in assay buffer (Hank's balanced salt solution with 25 mM HEPES pH 7.5) supplemented with 0.5 mM EGTA, followed by another wash in assay buffer. The cells were then resuspended in an assay buffer with 5 μg $ml^{-1}$ coelenterazine 400a (GoldBio) and placed in white 96-well plates (136101, Thermo Scientific) in a volume of 60 μl per well. 30 μl of ligands prepared at 3-times the final concentrations in assay buffer with 1.5 mM $CaCl_2$, 0.1% BSA, and 3% DMSO were added to plated cells (final concentrations of 0.5 mM $CaCl_2$, 0.33% BSA, and 1% DMSO). After 5 minutes of incubation, the emission at 410 and 515 nm were read using a SpectraMax iD5 plate reader with a 1-s integration time per well. The BRET ratios (GFP2/RLuc8 emission) were calculated and normalized to ligand-free control before further analysis. The efficacy and potency of the molecules were calculated by fitting the concentrations of molecules and the BRET ratios to a four-parameter logistic equation in Prism (Graphpad Software).

**Protein expression and purification**

CaSR-C1 and CaSR-C2 were overexpressed in *Spodoptera frugiperda* Sf9 cells using a Bac-to-Bac baculovirus expression system. Sf9 cells grown to a density of 3 x $10^6$ cells $ml^{-1}$ were co-infected with CaSR-C1 and CaSR-C2 baculoviruses for 48 h at 27°C. Cells were then harvested and stored at −80 °C. Purifications of CaSR in complex with compounds '54159 and '6218 followed a similar protocol. Cell pellets were thawed, resuspended, and lysed by nitrogen cavitation in the lysis buffer containing 20 mM HEPES pH 7.5, 150 mM NaCl, 10 mM $CaCl_2$, 10% glycerol, 10 mM L-Trp, protease inhibitors, benzonase, and 50 μM of a specific compound. The lysates were centrifuged at 1,000g for 10 min to remove nuclei and unlysed cells. The membranes

`

were harvested by centrifugation at 100,000g for 30 min and solubilized in the lysis buffer supplemented with 1% (w/v) Lauryl Maltose Neopentyl Glycol (LMNG, Anatrace) and 0.2% (w/v) cholesteryl hemisuccinate (CHS, Anatrace) for 3 h, followed by the centrifugation at 100,000g for 30 min. The supernatant was incubated with Strep-Tactin®XT 4Flow® resin (IBA) for overnight at 4°C. The resin was then loaded into a gravity column and washed with 10 column volumes of the washing buffer containing 20 mM HEPES 7.5, 150 mM NaCl, 10 mM $CaCl_2$, 5% glycerol, 40 μM L-Trp, and 50 μM compound, supplemented with 0.01% (w/v) LMNG and 0.002% (w/v) CHS, followed by a second wash with 10 column volumes of washing buffer with 0.001% (w/v) LMNG and 0.0002% (w/v) CHS. Proteins were eluted by Strep-Tactin®XT elution buffer (IBA) supplemented with 10 mM $CaCl_2$, 40 μM L-Trp, 50 μM compound, 0.00075% (w/v) LMNG, 0.00025% (w/v) GDN (CHS, Anatrace) and 0.00015% (w/v) CHS, and further purified by a Superose 6 column (Cytiva) using a buffer containing 20 mM HEPES 7.5, 150 mM NaCl, 10 mM $CaCl_2$, 40 μM L-Trp, 50 μM compound and 0.00075% (w/v) LMNG, 0.00025% (w/v) GDN and 0.00015% (w/v) CHS. The peak fractions were pooled and concentrated for cryo-EM studies.

**Cryo-EM data acquisition and data processing**

For cryo-EM imaging of the CaSR-'6218 complex, movies were collected using a Titan Krios G2 (Thermo Fisher Scientific) transmission electron microscope equipped with a Gatan K3 direct detector and a post-column energy filter with a 20 eV slit width. The microscope was operated at 300 kV, with a nominal magnification of 130,000x, resulting in a pixel size of 0.8677 Å. Movies were automatically recorded in counting mode using SerialEM (*58*) with a total exposure of 55 electrons·$Å^{-2}$ over 60 frames, and the defocus range was set from -0.5 to -1.5 μm. For cryo-EM imaging of the CaSR-'54159 complex, movies were collected using a Titan Krios G2 transmission electron microscope equipped with a Falcon 4i Direct Electron Detector and a post-column energy filter with a 20 eV slit width. The microscope was operated at 300 kV, with a nominal magnification of 165,000x, resulting in a pixel size of 0.75 Å. Movies were recorded in counting mode using

EPU 3.6 (Thermo Fisher Scientific) with a total exposure of 50 electrons·$Å^{-2}$ over 50 frames, and the defocus range was set from -0.5 to -1.5 μm.

For a detailed workflow of data processing, please refer to Extended Data Fig. 4. All data underwent processing using similar strategies using cryoSPARC 3.0 (*59*) and Relion 3 (*60*). Movies were imported into cryoSPARC and subjected to patch motion correction, followed by the contrast transfer function (CTF) estimation using patch CTF estimation. Micrographs with CTF estimations worse than 4 Å were excluded, resulting in a total of 11,926 micrographs for the CaSR-'6218 complex, and 17,625 micrographs for the CaSR-'54149 complexes, which were selected for further processing. Particles were autopicked, extracted from the micrographs, and subjected to 3-5 rounds of 2D classification. Particles classified into "good" classes were selected and subjected to iterative rounds of 3D ab initio reconstruction using multiple classes, followed by 3D heterogeneous refinement to remove particles from bad classes. For the early rounds of 3D classification, particles from "bad" classes were further classified by 2D classification and good particles were retained for subsequent heterogeneous refinement. The resulting high-quality particle projections were then imported into Relion, where they were subjected to C2 symmetry expansion, followed by 2-3 rounds of focused 3D classification (without applying symmetry) without alignment with a mask covering the two 7TMs of CaSR. Finally, the particles from one of the two best 3D classes with C1 symmetry were selected and imported to cryoSPARC for CTF refinement and local nonuniform refinement with a soft mask covering CRD–7TM and ECD-CRD to obtain high-resolution maps. The focused maps were used to generate composite maps for refinement.

**Model building and refinement**

The initial models of CaSR were built on the structure of the active-state CaSR (PDB ID: 7M3F) and manually docked into the cryo-EM maps in Chimera (*61*). The models were then subjected to iterative rounds of manual refinement in Coot (*62*) and automatic real-space refinement in

Phenix (*63*). The models for CRD–7TM and ECD-CRD regions were refined using the focused maps that cover these regions first and then combined for further refinement using the composite maps. The final models were analyzed and validated using MolProbity (*64*). The refinement statistics are shown in Extended Data Table 2. Structure figures were generated using ChimeraX (*65*).

**Animal studies**

Pharmacokinetics (PK) studies were performed on 10-weekold male CD1 mice by BIENTA Enamine Biology Services (Kiev, Ukraine). Briefly, the animals were randomly assigned to treatment groups for 9 time points (5, 15, 30, 60, 120, 240, 360, 480, and 1440 min) and fasted for 4 h before dosing with each PAM by subcutaneous (SC) route. At each time point post-injection, mice were injected IP with 2,2,2-tribromoethanol at the dose of 150 mg/kg prior to blood draws. All other animal studied were performed on 12-16 weeks old male C57/B6 mice (Jackson Laboratory; Bar Harbor, Maine, USA), approved by the Institutional Animal Care and Use Committee of the San Francisco Department of Veteran Affairs Medical Center (Protocol numbers: 2021–005 and 2021–016). For the latter studies, test compounds with specified doses were injected subcutaneously for 6 different time points (15, 30, 60, 120, 240, and 480 min), followed by isoflurane overdose before blood collections by cardiac puncture. Sera were prepared by centrifugation (2000xg) in microtainer (Becton Dickinson, SST 365967) and assayed for PTH levels by ELISA (Quidel, 60-2305) and total calcium using Alfa Wassermann ACE Axcel Vet Chemistry Analyzer.

***Ex vivo* parathyroid gland culture**

Mouse PTGs were isolated from 4-week-old male C57/B6 mice, dissected free of thyroid and surrounding fibrous tissues, and cultured to assess PTH secretion rate (ng/gland/hr) and $Ca^{2+}$ set-

`

point ($[Ca^{2+}]_e$ needed to suppress 50% of $PTH_{max}$) (*66, 67*). Briefly, PTGs were incubated sequentially with a series of DMEM media containing increasing concentrations of PAM at 0.75 mM calcium or containing increasing $[Ca^{2+}]_e$ with (50 or 500 nM) or without PAM to be tested. Intact PTH levels in culture media were assessed by ELISA and use to calculate the $EC_{50}$ or $Ca^{2+}$ set-points for each PAM.

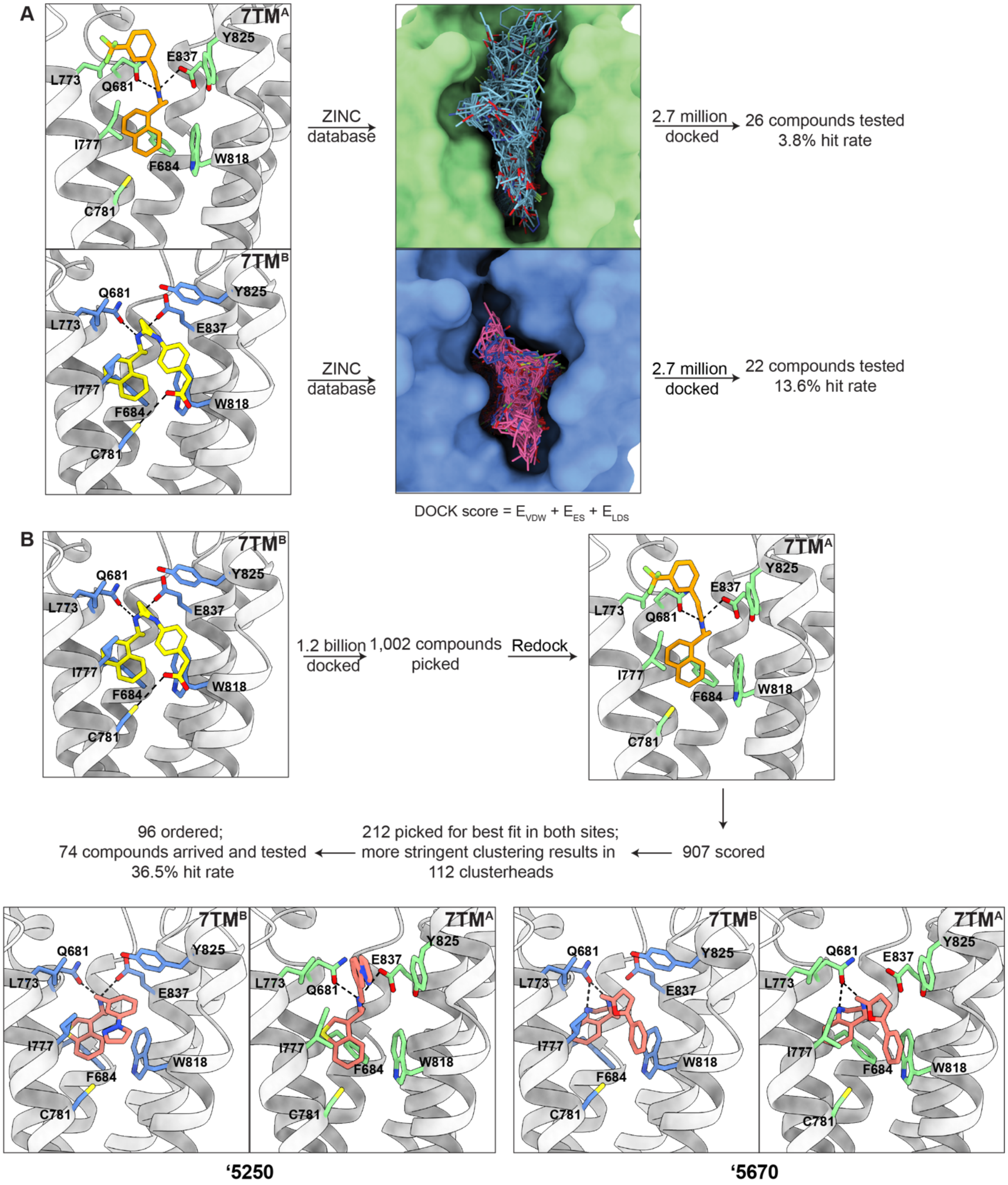
A
7TM^A
Y825
E837
L773
Q681
I777
F684
W818
C781
7TM^B
ZINC database
2.7 million docked
26 compounds tested
3.8% hit rate
22 compounds tested
13.6% hit rate
DOCK score = E_VDW + E_ES + E_LDS
B
1.2 billion docked
1,002 compounds picked
Redock
907 scored
212 picked for best fit in both sites;
more stringent clustering results in
112 clusterheads
96 ordered;
74 compounds arrived and tested
36.5% hit rate
'5250
'5670

`

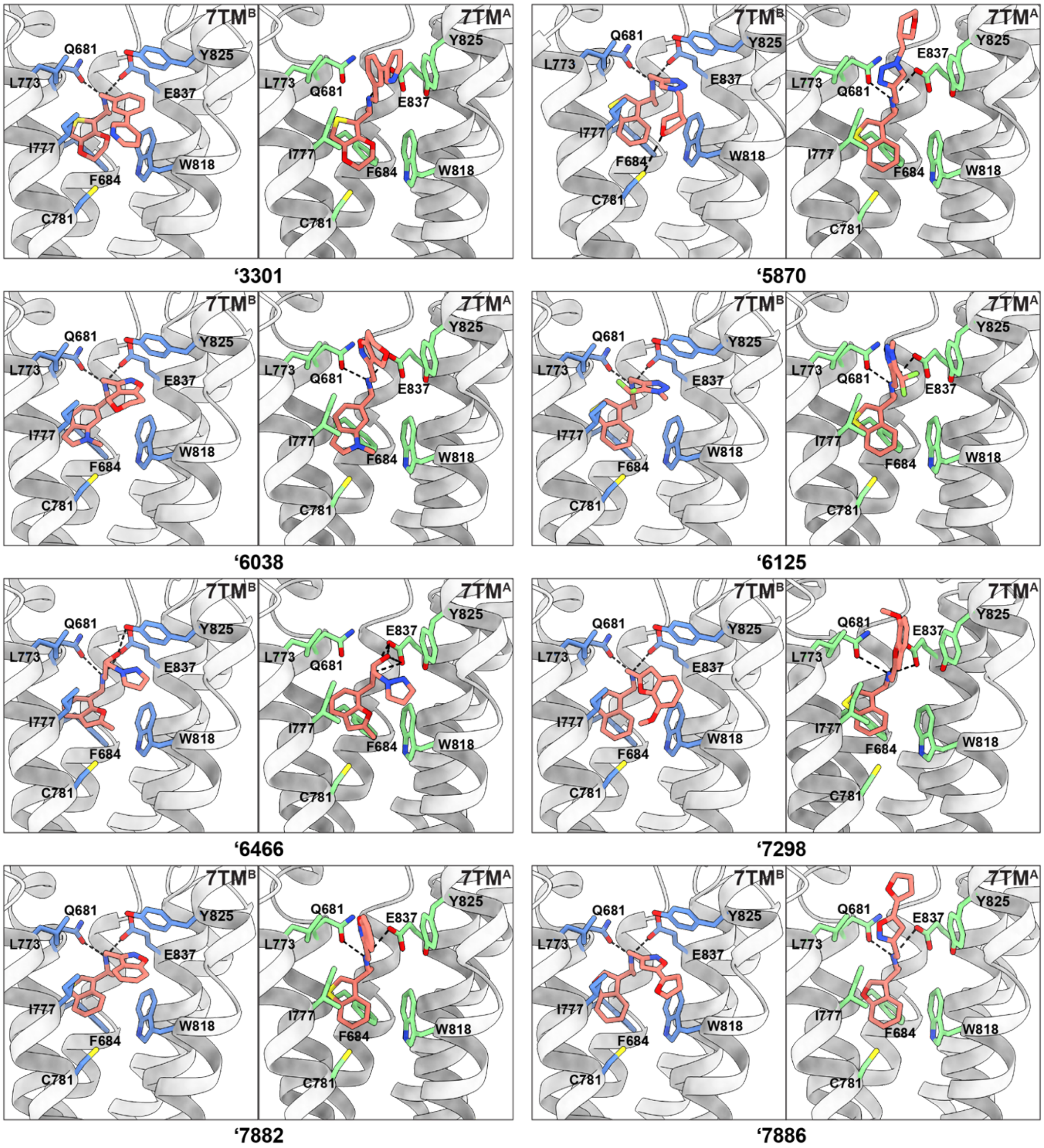

7TMB
7TMA
Q681
L773
Y825
E837
I777
F684
W818
C781
'3301
'5870
'6038
'6125
'6466
'7298
'7882
'7886

`

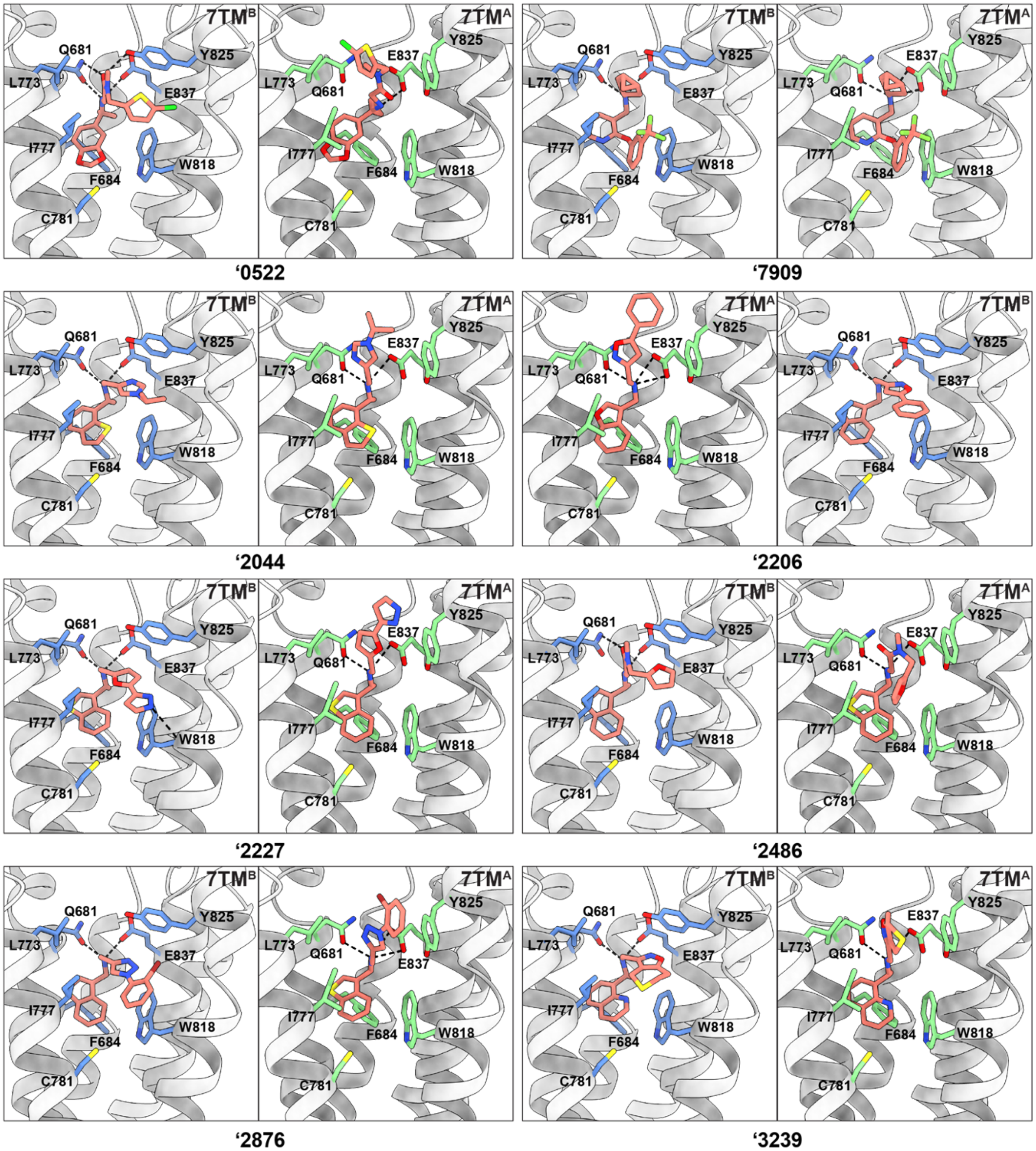
7TMB
7TMA
Q681
L773
Y825
E837
I777
F684
W818
C781
'0522
'7909
'2044
'2206
'2227
'2486
'2876
'3239

`

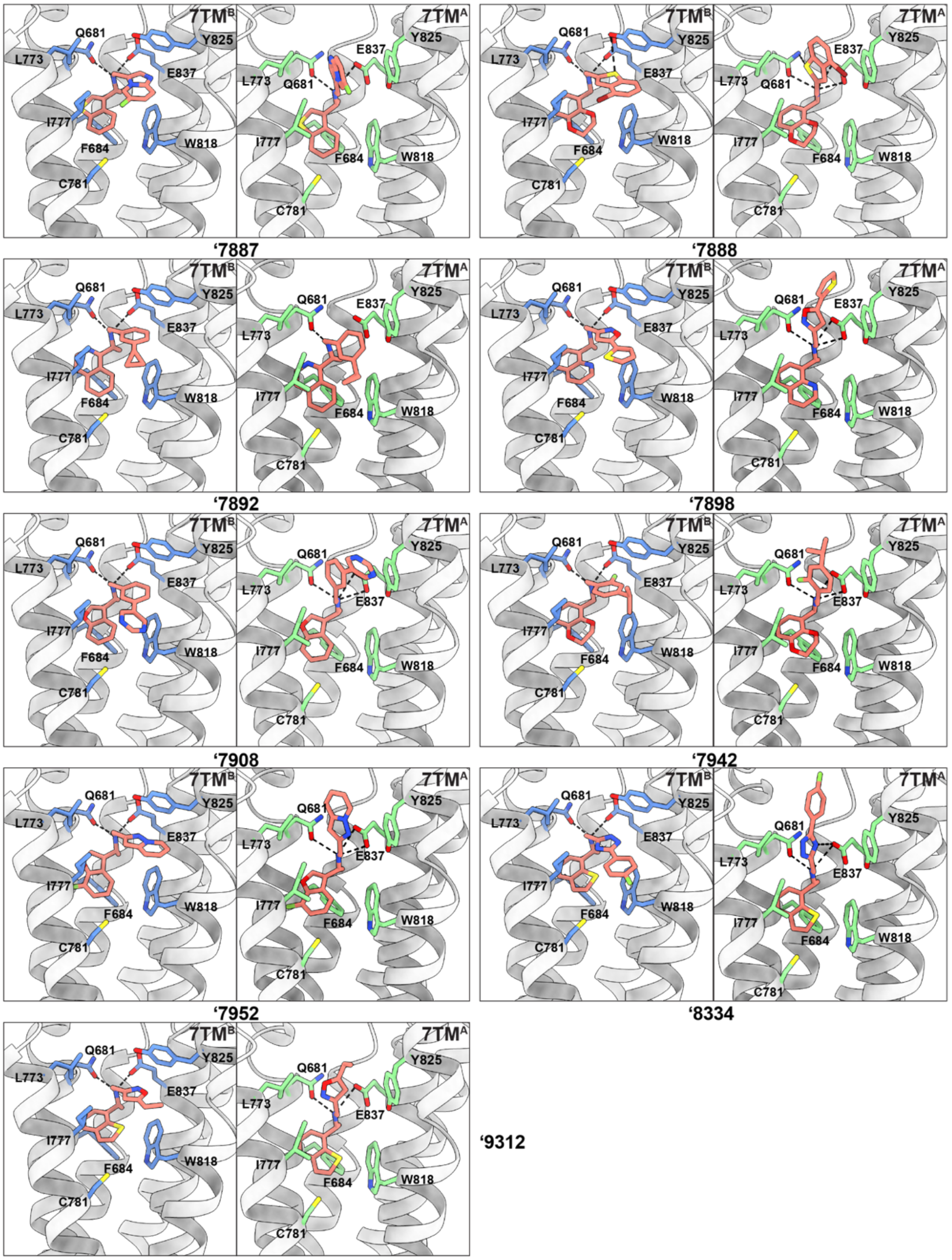


**Fig. S1: Docking workflow and the docked poses of the initial hits from the larger screen.** (**A**) Docking against 7TM$^B$ outperforms 7TM$^A$ in terms of higher hit rate when 2.7 million molecules were screened (13.6% versus 3.8%). (**B**) For the 1.2-billion docking, after docking against the 7TM$^B$ site, 1002 were picked after inspecting the top-scoring molecules. To reduce the number of candidate molecules, the 1,002 molecules were redocked against the 7TM$^A$ site. The 907 scored molecules were inspected again for best interactions with both binding sites, and further clustered for purchasing. Docked poses of the initial hits from large-scale docking campaign in 7TM$^A$ and 7TM$^B$ pockets are shown.

`

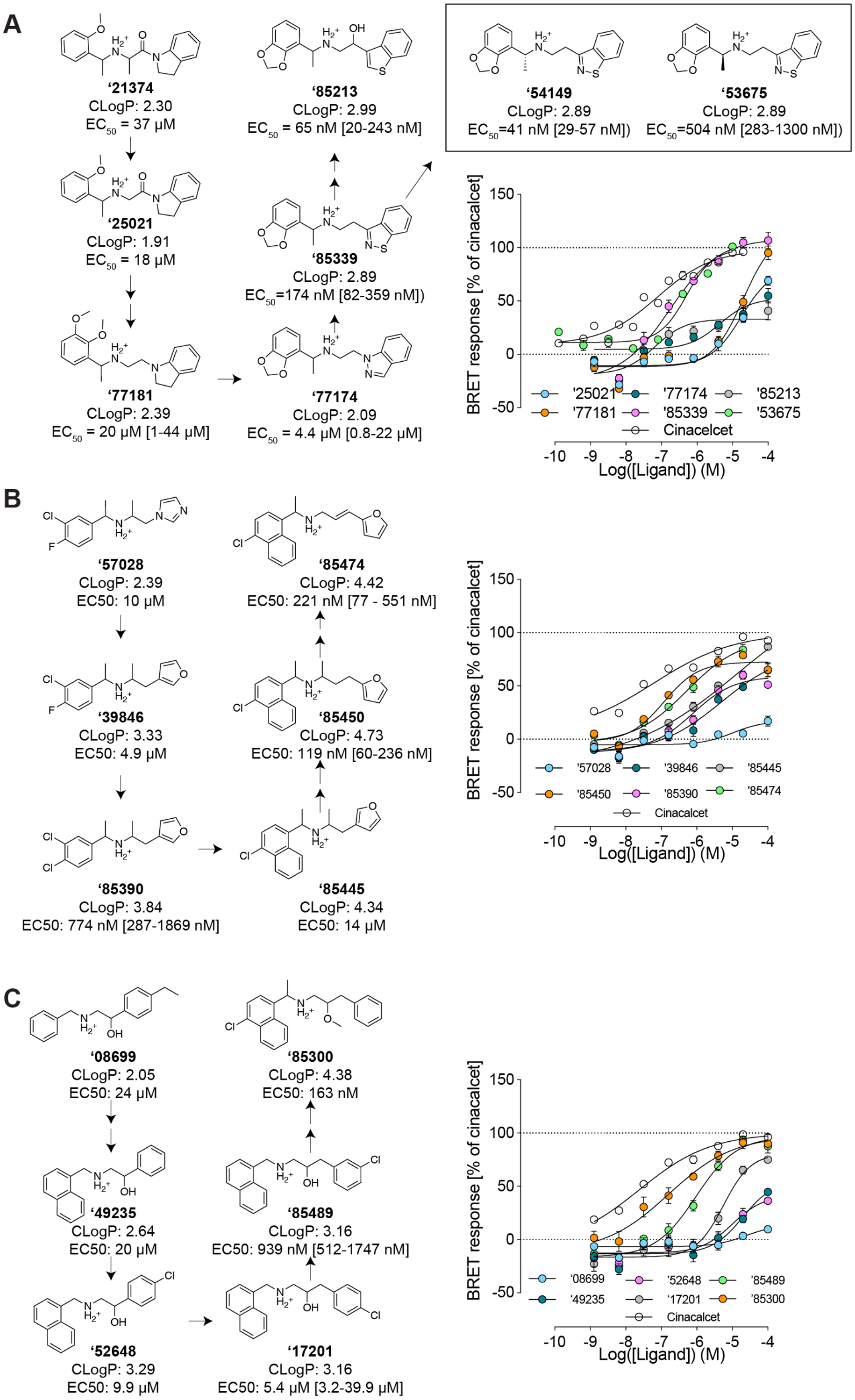


**Fig. S2: Structure activity relationships around the hits from "in-stock" screens**. (**A**) Additional SAR for optimization of compound **'21374.** (**B**) SAR for optimization of compound **'57028**. (**C**) SAR for optimization of compound **'08699**.

`

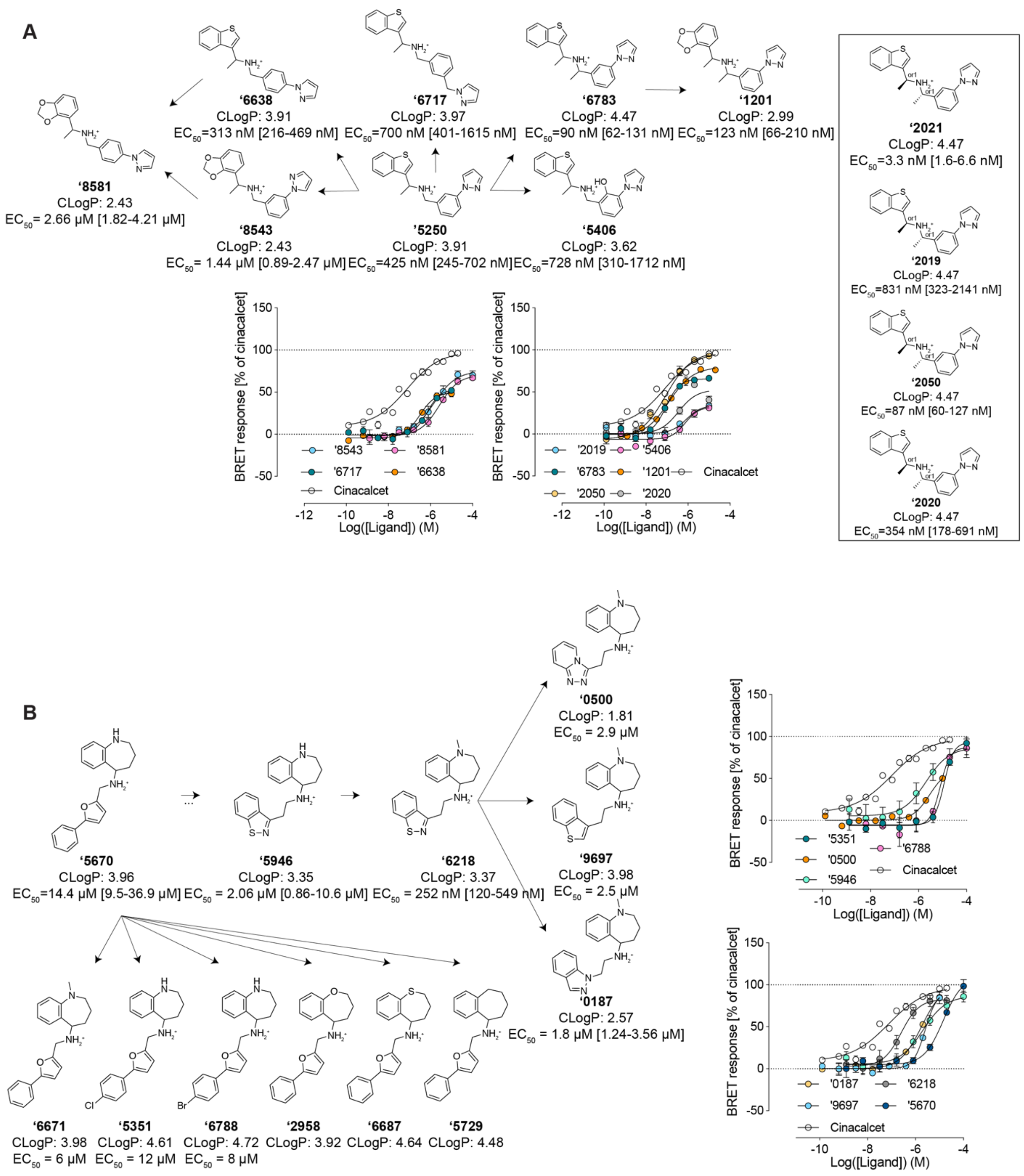
A
'6638
CLogP: 3.91
EC50=313 nM [216-469 nM]
'6717
CLogP: 3.97
EC50=700 nM [401-1615 nM]
'6783
CLogP: 4.47
EC50=90 nM [62-131 nM]
'1201
CLogP: 2.99
EC50=123 nM [66-210 nM]
'8581
CLogP: 2.43
EC50= 2.66 μM [1.82-4.21 μM]
'8543
CLogP: 2.43
EC50= 1.44 μM [0.89-2.47 μM]
'5250
CLogP: 3.91
EC50=425 nM [245-702 nM]
'5406
CLogP: 3.62
EC50=728 nM [310-1712 nM]
'2021
CLogP: 4.47
EC50=3.3 nM [1.6-6.6 nM]
'2019
CLogP: 4.47
EC50=831 nM [323-2141 nM]
'2050
CLogP: 4.47
EC50=87 nM [60-127 nM]
'2020
CLogP: 4.47
EC50=354 nM [178-691 nM]
BRET response [% of cinacalcet]
Log([Ligand]) (M)
'8543
'8581
'6717
'6638
Cinacalcet
'2019
'5406
'6783
'1201
Cinacalcet
'2050
'2020
B
'5670
CLogP: 3.96
EC50=14.4 μM [9.5-36.9 μM]
'5946
CLogP: 3.35
EC50 = 2.06 μM [0.86-10.6 μM]
'6218
CLogP: 3.37
EC50 = 252 nM [120-549 nM]
'0500
CLogP: 1.81
EC50 = 2.9 μM
'9697
CLogP: 3.98
EC50 = 2.5 μM
'0187
CLogP: 2.57
EC50 = 1.8 μM [1.24-3.56 μM]
'6671
CLogP: 3.98
EC50 = 6 μM
'5351
CLogP: 4.61
EC50 = 12 μM
'6788
CLogP: 4.72
EC50 = 8 μM
'2958
CLogP: 3.92
'6687
CLogP: 4.64
'5729
CLogP: 4.48
BRET response [% of cinacalcet]
Log([Ligand]) (M)
'5351
'6788
'0500
Cinacalcet
'5946
'0187
'6218
'9697
'5670
Cinacalcet

`

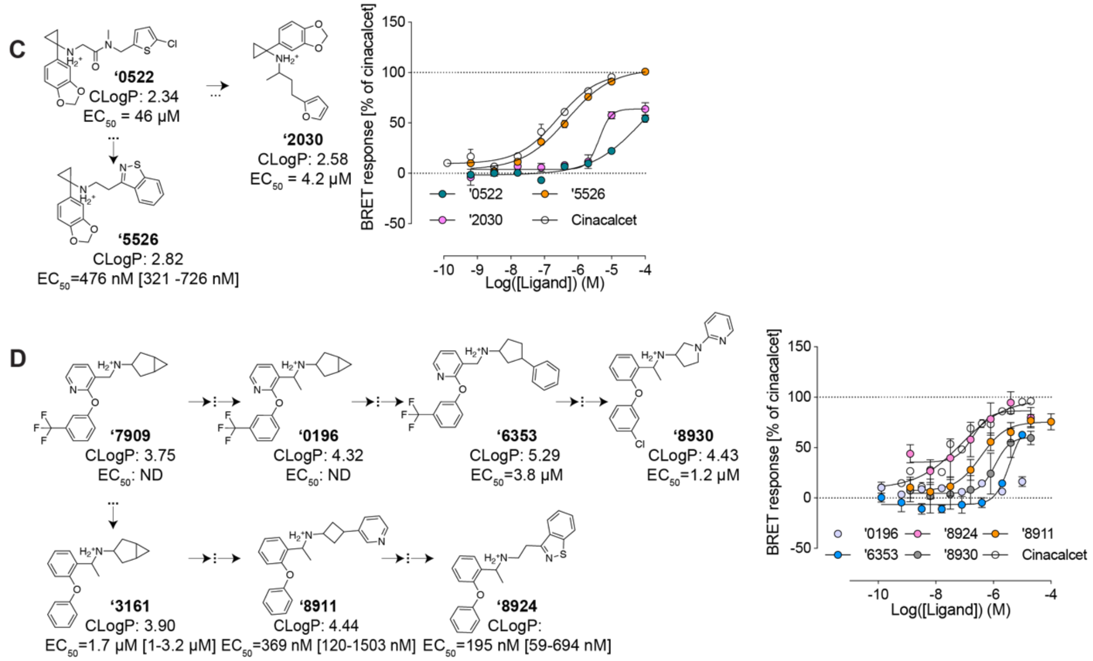


**Fig. S3. Additional structure activity relationships around the initial hits from large-scale docking campaign.** (**A**) Additional SAR for optimization of compound **'5250**. (**B**) Additional SAR for optimization of compound **'5670**. (**C**) SAR for optimization of compound **'0522**. (**D**) Additional SAR for optimization of compound **'7909**.

`

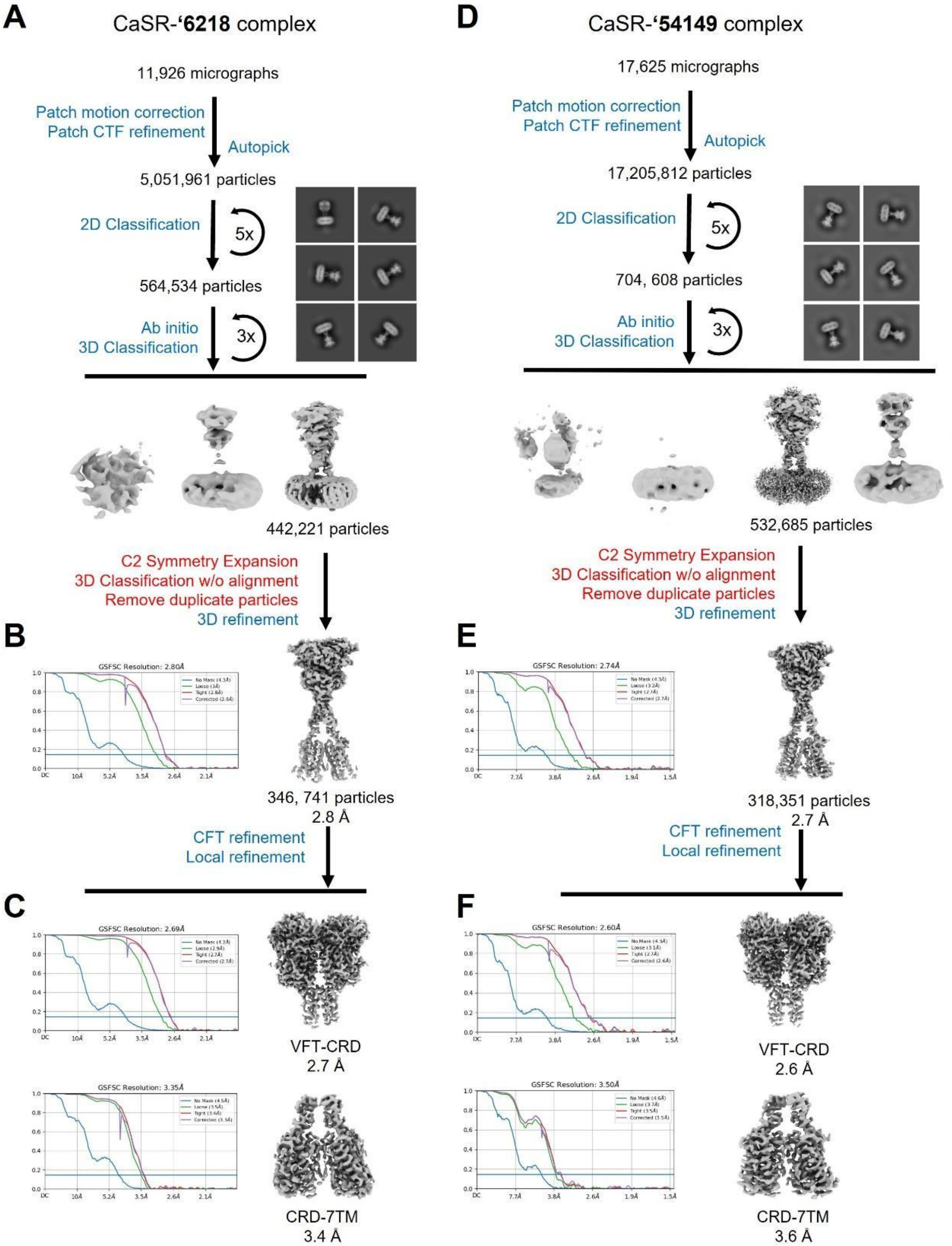


**Fig. S4: Cryo-EM processing workflow and reconstructions of the CaSR-'6218 and CaSR-'54149 complexes.** (**A**) Cryo-EM data processing workflow for CaSR-**'6218** complex. (**B**) The global map and (**C**) local maps of VFT–CRD, and CRD–7TM regions of CaSR-**'6218** complex with corresponding Fourier shell correlation (FSC) curves indicating nominal resolutions using the FSC = 0.143 criterion. (**D**) Cryo-EM data processing workflow for the CaSR-**'54149** complex. (**E**) The global map and (**F**) local maps of VFT–CRD, and CRD–7TM regions with FSC curves indicating nominal resolutions using the FSC = 0.143 criterion. Representative 2D averages are shown in square boxes (black) for each complex.

`

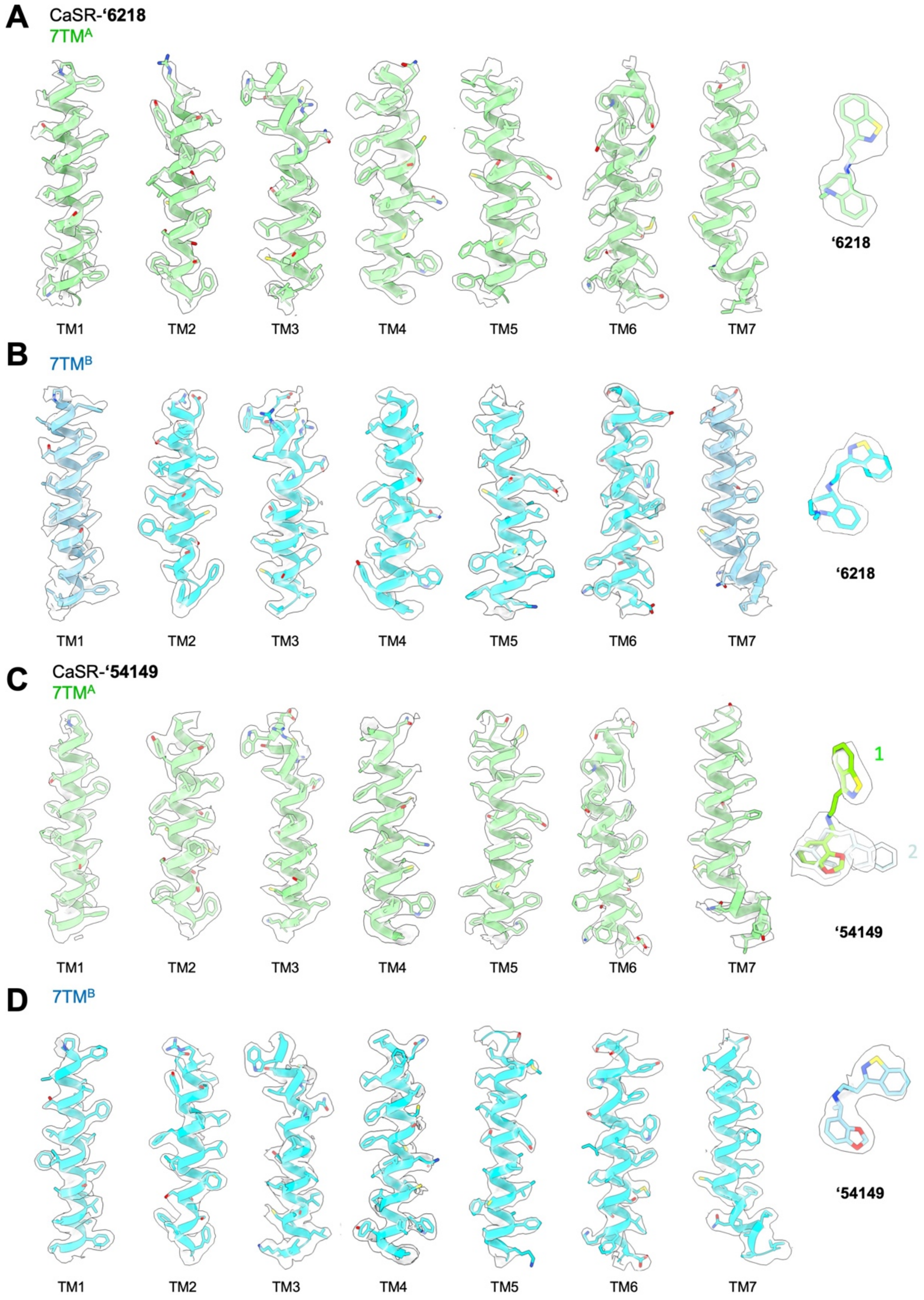


**Fig. S5: Agreement between cryo-EM density and model.** Models and their EM densities for the 7TM bundles and compound **'6218** in (**A**) 7TM$^{A}$ and (**B**) 7TM$^{B}$ of **CaSR-'6218** complex. Models and their EM densities for the 7TM bundles and compound **'54149** in (**C**) 7TM$^{A}$ and (**D**) 7TM$^{B}$ of CaSR-**'54149** complex. Models related to the 7TM$^{A}$ and 7TM$^{B}$ are colored in green and cyan, respectively. In (**C**), straight and folded-over conformations of **'54149** fitting into the density are shown in bright green and light cyan, respectively.

`

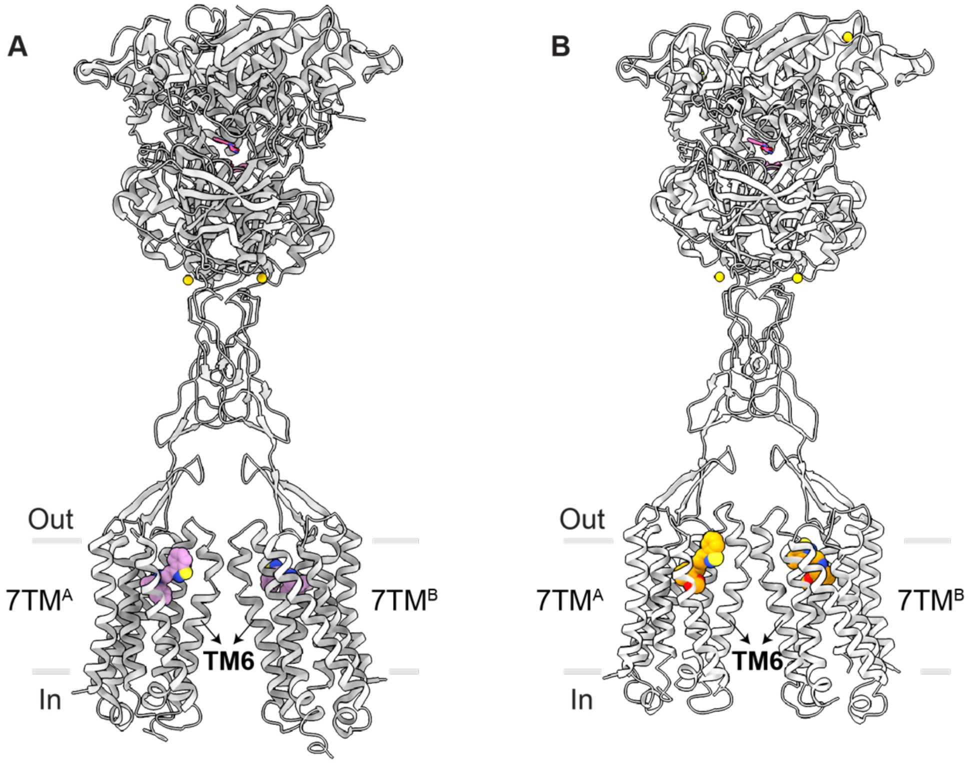


**Fig. S6: Asymmetric 7TMs configurations in CaSR complexes**. (**A**) Overall structure of **‘6218** bound CaSR. **‘6218** is in pink spheres. (**B**) Overall structure of **‘54149** bound CaSR. **‘54149** is in orange spheres. A higher sitting position of TM6 of 7TM$^A$ relative to 7TM$^B$ indicates an asymmetry arrangement of two 7TMs of CaSR.

`

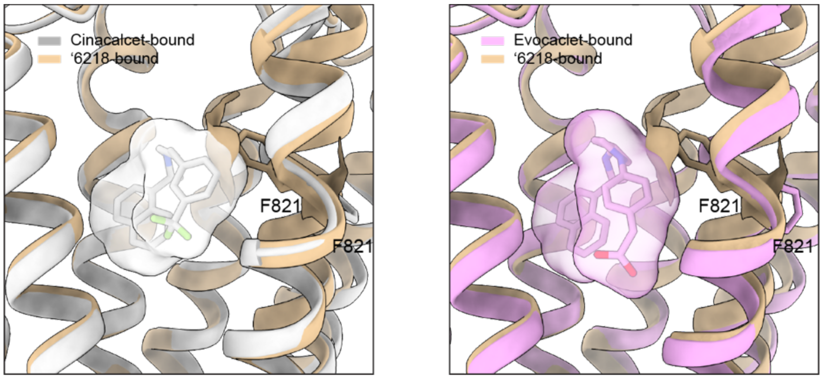


**Fig. S7: Structure comparison between the cinacalcet-bound CaSR or evocalcet-bound CaSR with ‘6218-bound CaSR in the 7TM$^B$ pocket.** Cinacalcet-bound CaSR is in silver, **‘6218-**bound CaSR (**‘6218** is hidden for clarity) is in tan. Evocalcet-bound CaSR is in pink. Residue F821’s side chain is shown. Evocalcet, cinacalcet and F821 from the ‘6218 structure are shown to demonstrate the clashes between them.

`

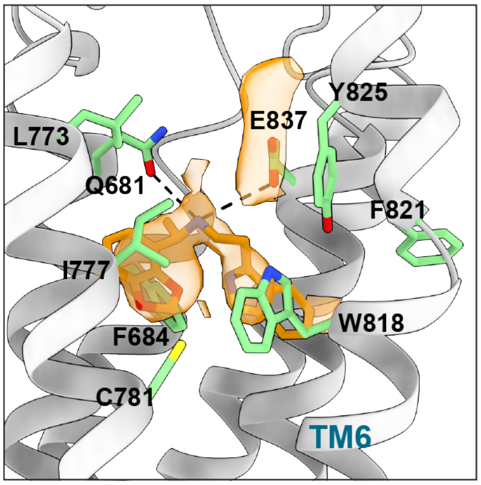


**Fig. S8: The "folded-over" pose of '54149 bound in the 7TM^A pocket.** Close-up view of "folded-over" conformation of **'54149** in the 7TM^A site, with EM density in the ligand-binding pocket shown in orange. Surrounding and key residues involved in the interaction are shown in green sticks.

`

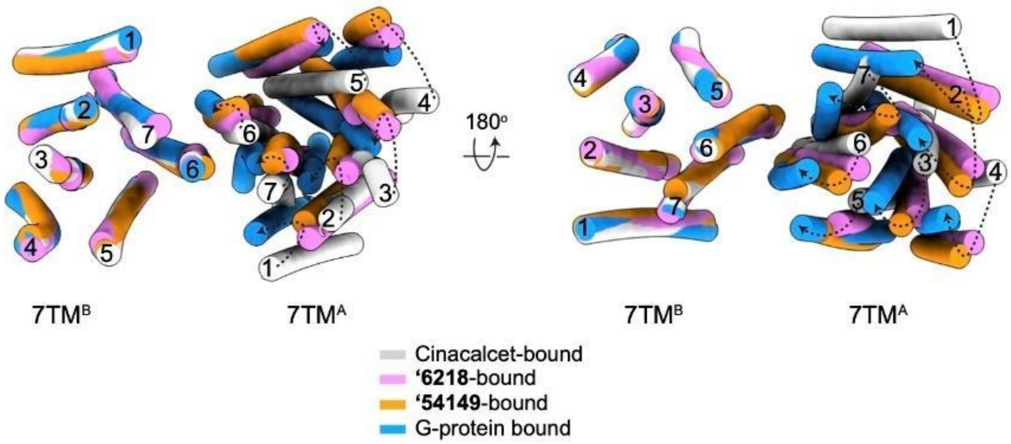


**Fig. S9: Comparison of overall arrangements of 7TMs in CaSR homodimer stabilized by different PAMs to G protein-bound active-state CaSR complex.** Overlay of the 7TMs of cinacalcet (grey, PDB: 7M3F),**’6218** (pink) and **‘54149** (orange)-bound CaSR, and $Gi_3$-bound CaSR (PDB: 8SZH) on the $7TM^B$ protomer. Top (left) and bottom (right) views are shown. The differences between the corresponding 7TM bundles in PAM-bound CaSR and $Gi_3$-bound CaSR are indicated by dashed arrows.

**Table S1. Potency for hits identified in initial CaSR docking screen and their Tanimoto coefficients (Tc) to known modulators.**

| Compound | Category | EC50 (µM) | Tc[b] | Nearest ChEMBL ligand[c] |
|---|---|---|---|---|
| 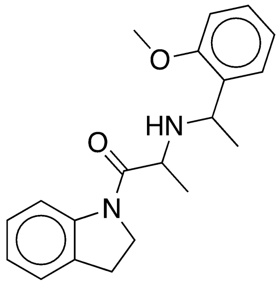 '21374 | In-stock initial hit | 37 | 0.28 | 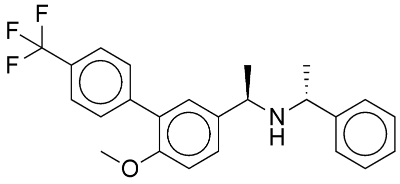 CHEMBL1224424 |
| 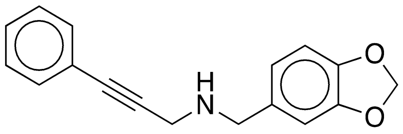 '74843 | In-stock initial hit | ND | 0.32 | 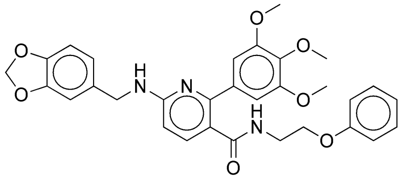 CHEMBL451383 |
| 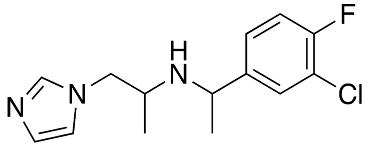 '57028 | In-stock initial hit | 10 | 0.28 | 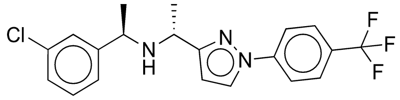 CHEMBL568485 |
| 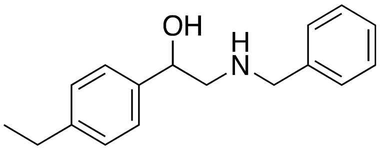 '08699 | In-stock initial hit | 24 | 0.34 | 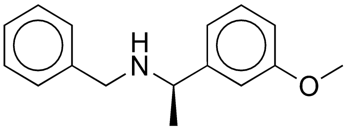 CHEMBL1224190 |
| 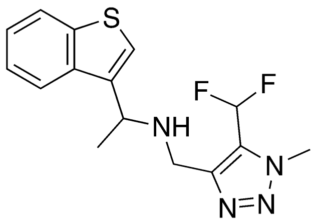 '6125 | LSD initial hit | 7.49 [4.82 – 17.95 ] | 0.3 | 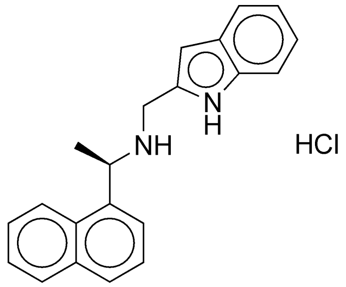 CHEMBL2092942 |
| 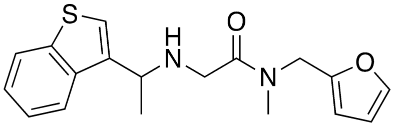 '2486 | LSD initial hit | 1.79 [1.02 – 3.93] | 0.25 | 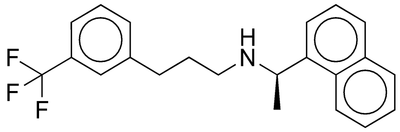 CHEMBL1201284 |

`

| | | | | |
|---|---|---|---|---|
| 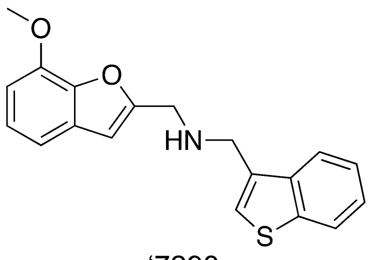 '7298 | LSD initial hit | 1.89 [1.19 – 2.95] | 0.27 | 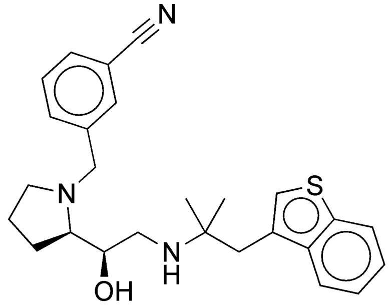 CHEMBL200041 |
| 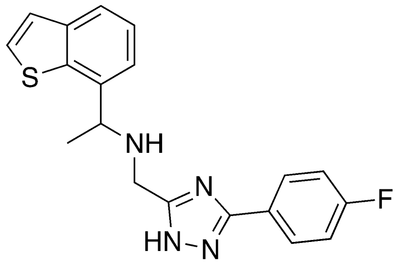 '8334 | LSD initial hit | 2.35 [1.21 – 7.98] | 0.31 | 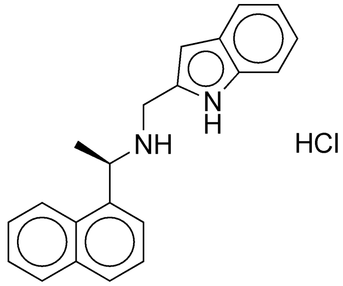 CHEMBL2092942 |
| 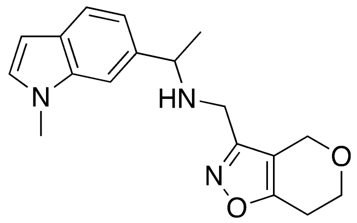 '6038 | LSD initial hit | ND | 0.25 | 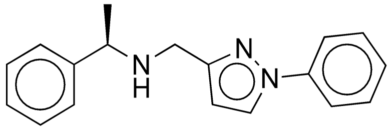 CHEMBL571048 |
| 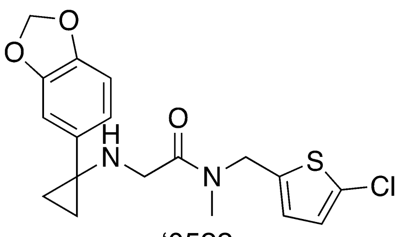 '0522 | LSD initial hit | 46.00 [10.73 – ??] | 0.25 | 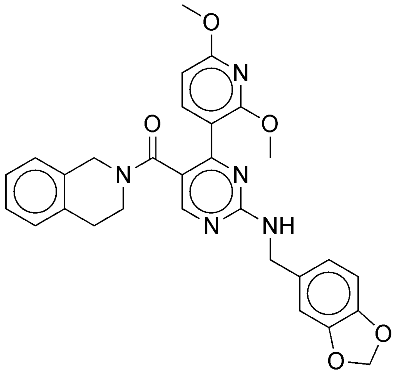 CHEMBL458515 |

`

| | | | | |
|---|---|---|---|---|
| 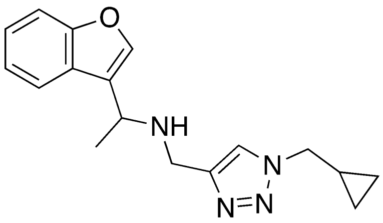<br>‘2700 | LSD initial hit | 3.08 [2.08 – 5.09] | 0.29 | 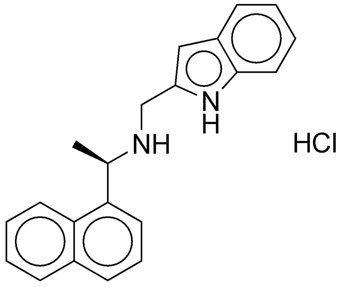<br>CHEMBL2092942 |
| 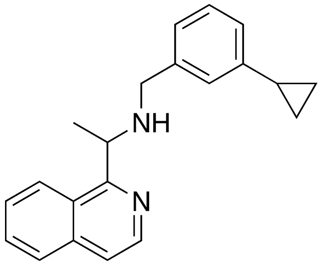<br>‘7892 | LSD initial hit | 6.34 [3.13 – 30.17 ] | 0.35 | 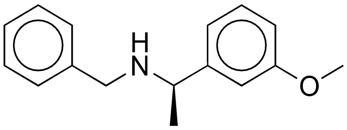<br>CHEMBL1224190 |
| 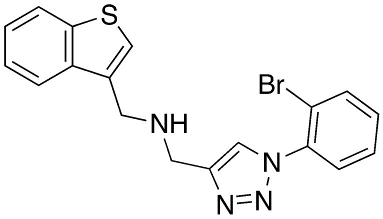<br>‘2876 | LSD initial hit | 4.82 [3.13 – 7.98] | 0.23 | 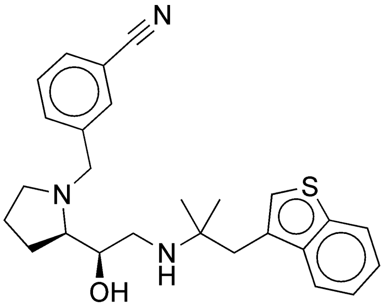<br>CHEMBL200041 |
| 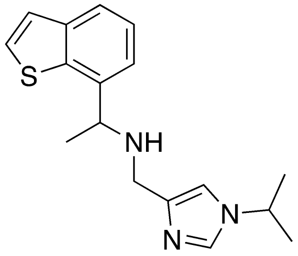<br>‘2044 | LSD initial hit | 4.41 [2.85 – 9.14] | 0.33 | 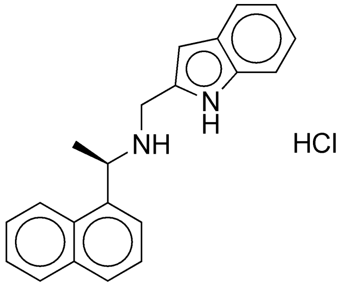<br>CHEMBL2092942 |
| 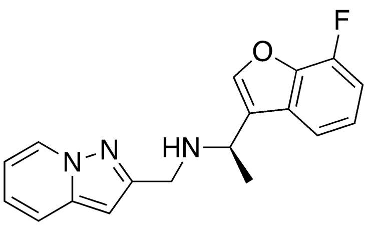<br>‘7952 | LSD initial hit | 0.273 [0.181 – 0.413 ] | 0.35 | 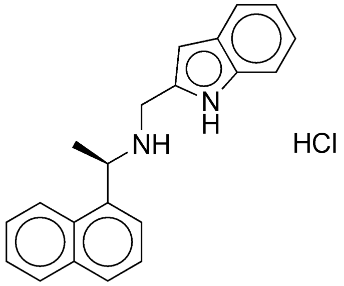<br>CHEMBL2092942 |

`

| | | | | |
|---|---|---|---|---|
| 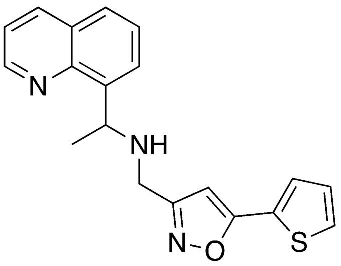 '7898 | LSD initial hit | 5.84 [2.94 – 32.49 ] | 0.35 | 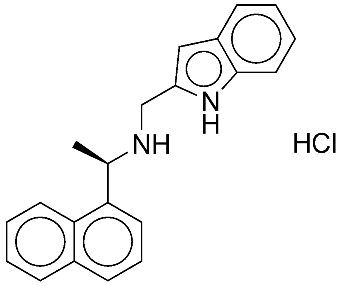 CHEMBL2092942 |
| 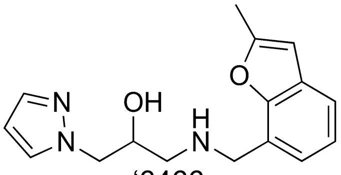 '6466 | LSD initial hit | ND | 0.26 | 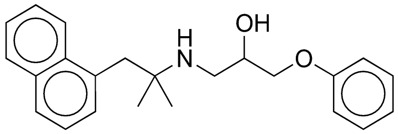 CHEMBL488736 |
| 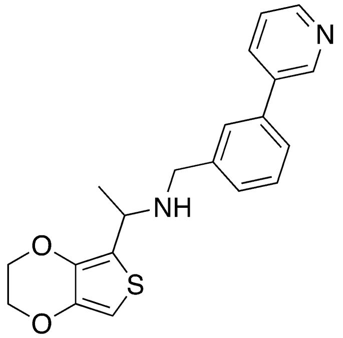 '3301 | LSD initial hit | 4.83 [2.46 – 65.73 ] | 0.28 | 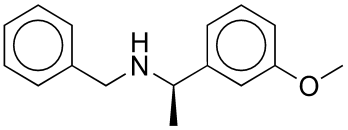 CHEMBL1224190 |
| 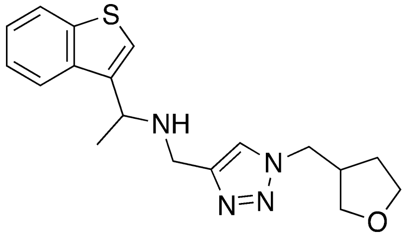 '5870 | LSD initial hit | 0.893 [0.662 – 1.225 ] | 0.27 | 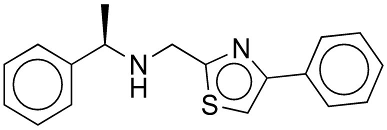 CHEMBL566524 |

`

| | | | | |
|---|---|---|---|---|
| 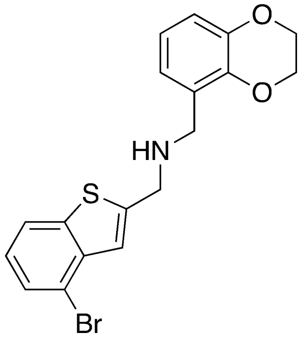 '7888 | LSD initial hit | 2.29 [1.86 – 2.92] | 0.18 | 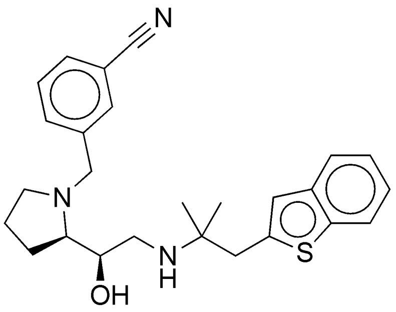 CHEMBL200312 |
| 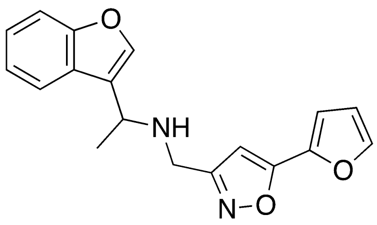 '7886 | LSD initial hit | 0.895 [0.464 – 3.265 ] | 0.34 | 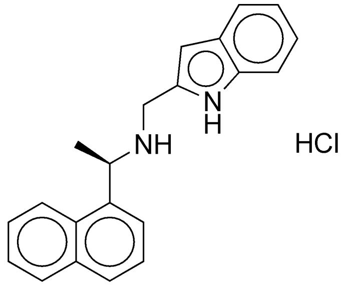 CHEMBL2092942 |
| 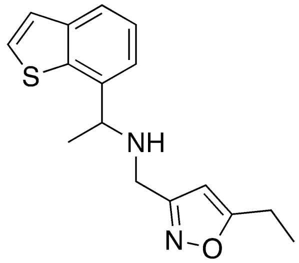 '9312 | LSD initial hit | 2.79 [1.74 – 5.01] | 0.34 | 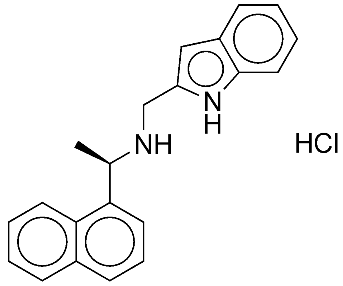 CHEMBL2092942 |
| 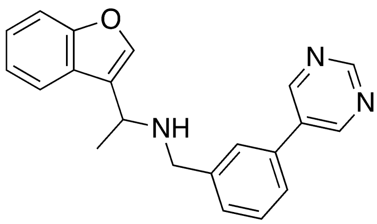 '7908 | LSD initial hit | 0.341 [0.207 – 0.556 ] | 0.34 | 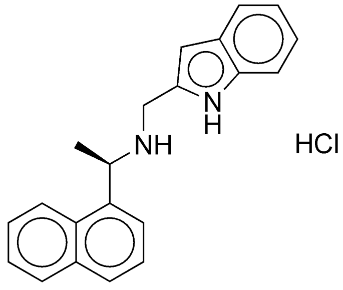 CHEMBL2092942 |

`

| | | | | |
|---|---|---|---|---|
| 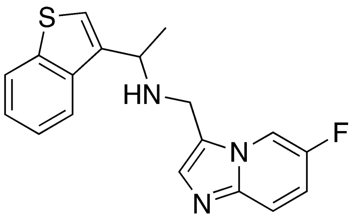 ‘7887 | LSD initial hit | 1.40 [0.69 – 4.36] | 0.29 | 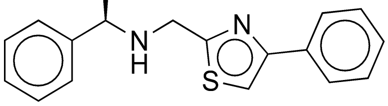 CHEMBL566524 |
| 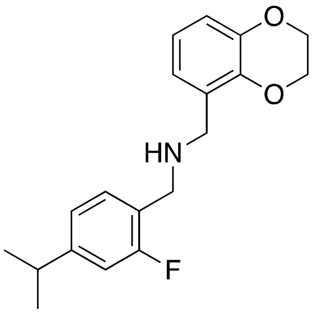 ‘7942 | LSD initial hit | ND | 0.25 | 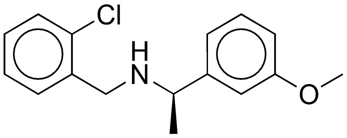 CHEMBL1224191 |
| 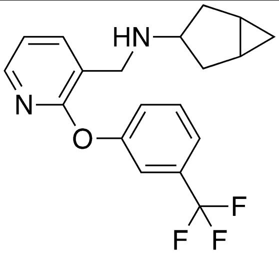 ‘7909 | LSD initial hit | ND | 0.28 | 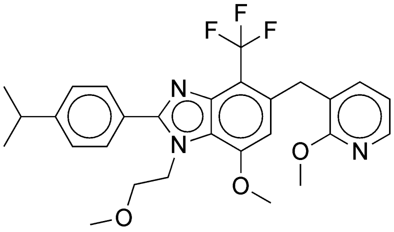 CHEMBL1223712 |
| 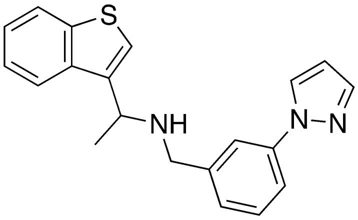 ‘5250 | LSD initial hit | 0.415 [0.245 – 0.790] | 0.32 | 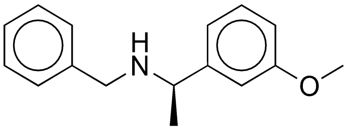 CHEMBL1224190 |
| 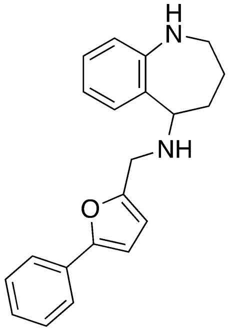 ‘5670 | LSD initial hit | 14.4 [9.5 – 36.9] | 0.22 | 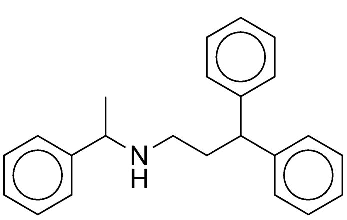 CHEMBL254832 |

`

| | | | | |
|---|---|---|---|---|
| 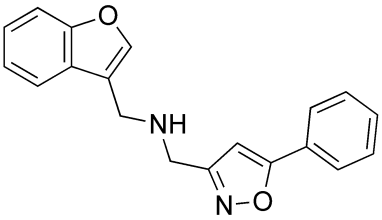 '2206 | LSD initial hit | ND | 0.24 | 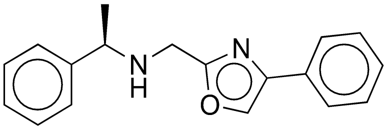 CHEMBL572172 |
| 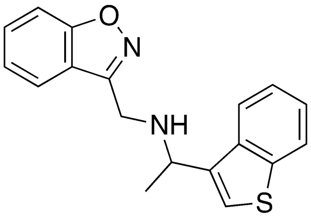 '7882 | LSD initial hit | 1.01 [0.61 – 2.39] | 0.31 | 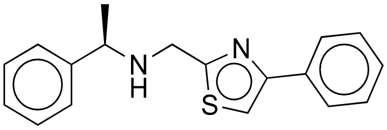 CHEMBL566524 |
| 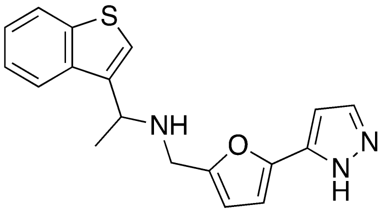 '2227 | LSD initial hit | 0.736 [0.452 – 1.623] | 0.28 | 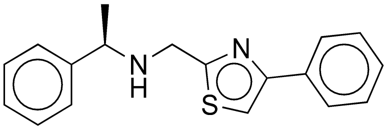 CHEMBL566524 |

`

**Table S2. Cryo-EM data collection, refinement and validation statistics**

| | CaSR-‘6218 | CaSR-‘54149 |
|---|---|---|
| **Data collection and processing** | | |
| Magnification (kx) | 130 | 165 |
| Voltage (kV) | 300 | 300 |
| Defocus (µm) | -0.5 to -1.5 | -0.5 to -1.5 |
| Pixel size (Å) | 0.8677 | 0.7500 |
| Total dose ($e^-$ / $Å^2$) | 55 | 50 |
| Symmetry imposed | C1 | C1 |
| Number of micrographs used | 11,926 | 17,625 |
| Number of initial particles picked | 5,051,961 | 1,720,812 |
| Number of final particles refined | 346,741 | 318,351 |
| Map resolution (Å) | 2.8 (global)<br>2.7 (VFT–CRD)<br>3.4 (CRD–7TM) | 2.7 (global)<br>2.6 (VFT–CRD)<br>3.6 (CRD–7TM) |
| FSC threshold (Å) | 0.143 | 0.143 |
| **Refinement** | | |
| Model composition | | |
| Non-hydrogen atoms | 12251 | 12243 |
| Ligand | 18 | 18 |
| Water | 0 | 0 |
| RMSD | | |
| bond length ($Å^2$) | 0.006 | 0.003 |
| bond angle (°) | 0.723 | 0.520 |
| Validation | | |
| MolProbity Score | 1.62 | 1.47 |
| Clash score | 5.62 | 5.09 |
| Ramachandran plot (%) | | |
| Favored | 96.07 | 96.78 |
| Allowed | 3.93 | 3.22 |
| Disallowed | 0 | 0 |

### References:

1. E. M. Brown *et al.*, Cloning and characterization of an extracellular Ca(2+)-sensing receptor from bovine parathyroid. *Nature* **366**, 575-580 (1993).
2. K. Leach *et al.*, International Union of Basic and Clinical Pharmacology. CVIII. Calcium-Sensing Receptor Nomenclature, Pharmacology, and Function. *Pharmacol Rev* **72**, 558-604 (2020).
3. F. M. Hannan, E. Kallay, W. Chang, M. L. Brandi, R. V. Thakker, The calcium-sensing receptor in physiology and in calcitropic and noncalcitropic diseases. *Nat Rev Endocrinol* **15**, 33-51 (2018).
4. F. M. Hannan, R. V. Thakker, Calcium-sensing receptor (CaSR) mutations and disorders of calcium, electrolyte and water metabolism. *Best Pract Res Clin Endocrinol Metab* **27**, 359-371 (2013).
5. M. R. Pollak *et al.*, Autosomal dominant hypocalcaemia caused by a Ca(2+)-sensing receptor gene mutation. *Nat Genet* **8**, 303-307 (1994).
6. F. M. Hannan *et al.*, Identification of 70 calcium-sensing receptor mutations in hyper- and hypo-calcaemic patients: evidence for clustering of extracellular domain mutations at calcium-binding sites. *Hum Mol Genet* **21**, 2768-2778 (2012).
7. S. H. Pearce *et al.*, A familial syndrome of hypocalcemia with hypercalciuria due to mutations in the calcium-sensing receptor. *N Engl J Med* **335**, 1115-1122 (1996).
8. J. Patel, M. B. Bridgeman, Etelcalcetide (Parsabiv) for Secondary Hyperparathyroidism in Adults With Chronic Kidney Disease on Hemodialysis. *P T* **43**, 396-399 (2018).
9. L. Pereira, C. Meng, D. Marques, J. M. Frazao, Old and new calcimimetics for treatment of secondary hyperparathyroidism: impact on biochemical and relevant clinical outcomes. *Clin Kidney J* **11**, 80-88 (2018).
10. T. C. Sauter *et al.*, Calcium Disorders in the Emergency Department: Independent Risk Factors for Mortality. *PLoS One* **10**, e0132788 (2015).
11. Z. Zhang, X. Xu, H. Ni, H. Deng, Predictive value of ionized calcium in critically ill patients: an analysis of a large clinical database MIMIC II. *PLoS One* **9**, e95204 (2014).
12. M. Egi *et al.*, Ionized calcium concentration and outcome in critical illness. *Crit Care Med* **39**, 314-321 (2011).
13. T. Steele, R. Kolamunnage-Dona, C. Downey, C. H. Toh, I. Welters, Assessment and clinical course of hypocalcemia in critical illness. *Crit Care* **17**, R106 (2013).
14. A. Husain, R. J. Simpson, Jr., G. Joodi, Serum Calcium and Risk of Sudden Cardiac Arrest in the General Population. *Mayo Clin Proc* **93**, 392 (2018).
15. R. Nardone, F. Brigo, E. Trinka, Acute Symptomatic Seizures Caused by Electrolyte Disturbances. *J Clin Neurol* **12**, 21-33 (2016).
16. T. B. Drueke, Cell biology of parathyroid gland hyperplasia in chronic renal failure. *J Am Soc Nephrol* **11**, 1141-1152 (2000).
17. J. C. Bureo *et al.*, Prevalence of secondary hyperparathyroidism in patients with stage 3 and 4 chronic kidney disease seen in internal medicine. *Endocrinol Nutr* **62**, 300-305 (2015).
18. A. Levin *et al.*, Prevalence of abnormal serum vitamin D, PTH, calcium, and phosphorus in patients with chronic kidney disease: results of the study to evaluate early kidney disease. *Kidney Int* **71**, 31-38 (2007).
19. D. L. Andress *et al.*, Management of secondary hyperparathyroidism in stages 3 and 4 chronic kidney disease. *Endocr Pract* **14**, 18-27 (2008).
20. C. P. Kovesdy, Epidemiology of chronic kidney disease: an update 2022. *Kidney Int Suppl (2011)* **12**, 7-11 (2022).

`


21. J. P. Pin, T. Galvez, L. Prezeau, Evolution, structure, and activation mechanism of family 3/C G-protein-coupled receptors. *Pharmacol Ther* **98**, 325-354 (2003).
22. Y. Gao *et al.*, Asymmetric activation of the calcium-sensing receptor homodimer. *Nature* **595**, 455-459 (2021).
23. A. B. Seven *et al.*, G-protein activation by a metabotropic glutamate receptor. *Nature* **595**, 450-454 (2021).
24. M. M. Papasergi-Scott *et al.*, Structures of metabotropic GABA(B) receptor. *Nature* **584**, 310-314 (2020).
25. J. Lyu *et al.*, Ultra-large library docking for discovering new chemotypes. *Nature* **566**, 224-229 (2019).
26. C. Gorgulla *et al.*, An open-source drug discovery platform enables ultra-large virtual screens. *Nature* **580**, 663-668 (2020).
27. R. M. Stein *et al.*, Virtual discovery of melatonin receptor ligands to modulate circadian rhythms. *Nature* **579**, 609-614 (2020).
28. A. Alon *et al.*, Structures of the sigma(2) receptor enable docking for bioactive ligand discovery. *Nature* **600**, 759-764 (2021).
29. A. A. Sadybekov *et al.*, Synthon-based ligand discovery in virtual libraries of over 11 billion compounds. *Nature* **601**, 452-459 (2022).
30. E. A. Fink *et al.*, Structure-based discovery of nonopioid analgesics acting through the alpha(2A)-adrenergic receptor. *Science* **377**, eabn7065 (2022).
31. I. Singh *et al.*, Structure-based discovery of conformationally selective inhibitors of the serotonin transporter. *Cell* **186**, 2160-2175 e2117 (2023).
32. R. G. Coleman, M. Carchia, T. Sterling, J. J. Irwin, B. K. Shoichet, Ligand pose and orientational sampling in molecular docking. *PLoS One* **8**, e75992 (2013).
33. E. C. Meng, B. K. Shoichet, I. D. Kuntz, Automated Docking with Grid-Based Energy Evaluation. *J Comput Chem* **13**, 505-524 (1992).
34. K. A. Sharp, R. A. Friedman, V. Misra, J. Hecht, B. Honig, Salt effects on polyelectrolyte-ligand binding: comparison of Poisson-Boltzmann, and limiting law/counterion binding models. *Biopolymers* **36**, 245-262 (1995).
35. K. Gallagher, K. Sharp, Electrostatic contributions to heat capacity changes of DNA-ligand binding. *Biophys J* **75**, 769-776 (1998).
36. M. M. Mysinger, B. K. Shoichet, Rapid context-dependent ligand desolvation in molecular docking. *J Chem Inf Model* **50**, 1561-1573 (2010).
37. S. Gu, M. S. Smith, Y. Yang, J. J. Irwin, B. K. Shoichet, Ligand Strain Energy in Large Library Docking. *J Chem Inf Model* **61**, 4331-4341 (2021).
38. R. H. J. Olsen *et al.*, TRUPATH, an open-source biosensor platform for interrogating the GPCR transducerome. *Nat Chem Biol* **16**, 841-849 (2020).
39. B. I. Tingle *et al.*, ZINC-22 horizontal line A Free Multi-Billion-Scale Database of Tangible Compounds for Ligand Discovery. *J Chem Inf Model* **63**, 1166-1176 (2023).
40. J. Lyu, J. J. Irwin, B. K. Shoichet, Modeling the expansion of virtual screening libraries. *Nat Chem Biol* **19**, 712-718 (2023).
41. K. Leach *et al.*, Towards a structural understanding of allosteric drugs at the human calcium-sensing receptor. *Cell Res* **26**, 574-592 (2016).
42. A. N. Keller *et al.*, Identification of Global and Ligand-Specific Calcium Sensing Receptor Activation Mechanisms. *Mol Pharmacol* **93**, 619-630 (2018).
43. F. He *et al.*, Allosteric modulation and G-protein selectivity of the $Ca^{2+}$-sensing receptor. *Nature*, (2024). Feb 7. doi: 10.1038/s41586-024-07055-2. Epub ahead of print. PMID: 38326620.
44. S. Lin *et al.*, Structures of G(i)-bound metabotropic glutamate receptors mGlu2 and mGlu4. *Nature* **594**, 583-588 (2021).

45. E. M. Brown, Clinical lessons from the calcium-sensing receptor. *Nat Clin Pract Endocrinol Metab* **3**, 122-133 (2007).
46. G. A. Block *et al.*, Effect of Etelcalcetide vs Cinacalcet on Serum Parathyroid Hormone in Patients Receiving Hemodialysis With Secondary Hyperparathyroidism: A Randomized Clinical Trial. *JAMA* **317**, 156-164 (2017).
47. S. A. Jamal, P. D. Miller, Secondary and tertiary hyperparathyroidism. *J Clin Densitom* **16**, 64-68 (2013).
48. M. Rodriguez, E. Nemeth, D. Martin, The calcium-sensing receptor: a key factor in the pathogenesis of secondary hyperparathyroidism. *Am J Physiol Renal Physiol* **288**, F253-264 (2005).
49. P. P. Centeno *et al.*, Phosphate acts directly on the calcium-sensing receptor to stimulate parathyroid hormone secretion. *Nat Commun* **10**, 4693 (2019).
50. J. Gogusev *et al.*, Depressed expression of calcium receptor in parathyroid gland tissue of patients with hyperparathyroidism. *Kidney Int* **51**, 328-336 (1997).
51. G. S. Schmidt, T. D. Weaver, T. D. Hoang, M. K. M. Shakir, Severe Symptomatic Hypocalcemia, complicating cardiac arrhythmia following Cinacalcet (Sensipar(TM)) administration: A Case Report. *Clin Case Rep* **9**, e04876 (2021).
52. G. A. Block *et al.*, Cinacalcet for secondary hyperparathyroidism in patients receiving hemodialysis. *N Engl J Med* **350**, 1516-1525 (2004).
53. J. M. Word, S. C. Lovell, J. S. Richardson, D. C. Richardson, Asparagine and glutamine: using hydrogen atom contacts in the choice of side-chain amide orientation. *J Mol Biol* **285**, 1735-1747 (1999).
54. G. M. Sastry, M. Adzhigirey, T. Day, R. Annabhimoju, W. Sherman, Protein and ligand preparation: parameters, protocols, and influence on virtual screening enrichments. *J Comput Aided Mol Des* **27**, 221-234 (2013).
55. K. A. Sharp, Polyelectrolyte Electrostatics - Salt Dependence, Entropic, and Enthalpic Contributions to Free-Energy in the Nonlinear Poisson-Boltzmann Model. *Biopolymers* **36**, 227-243 (1995).
56. R. M. Stein *et al.*, Property-Unmatched Decoys in Docking Benchmarks. *J Chem Inf Model* **61**, 699-714 (2021).
57. A. V. Fassio *et al.*, Prioritizing Virtual Screening with Interpretable Interaction Fingerprints. *J Chem Inf Model* **62**, 4300-4318 (2022).
58. D. N. Mastronarde, Automated electron microscope tomography using robust prediction of specimen movements. *J Struct Biol* **152**, 36-51 (2005).
59. A. Punjani, J. L. Rubinstein, D. J. Fleet, M. A. Brubaker, cryoSPARC: algorithms for rapid unsupervised cryo-EM structure determination. *Nat Methods* **14**, 290-296 (2017).
60. J. Zivanov *et al.*, New tools for automated high-resolution cryo-EM structure determination in RELION-3. *Elife* **7**, (2018).
61. E. F. Pettersen *et al.*, UCSF Chimera--a visualization system for exploratory research and analysis. *J Comput Chem* **25**, 1605-1612 (2004).
62. P. Emsley, B. Lohkamp, W. G. Scott, K. Cowtan, Features and development of Coot. *Acta Crystallogr D Biol Crystallogr* **66**, 486-501 (2010).
63. D. Liebschner *et al.*, Macromolecular structure determination using X-rays, neutrons and electrons: recent developments in Phenix. *Acta Crystallogr D Struct Biol* **75**, 861-877 (2019).
64. V. B. Chen *et al.*, MolProbity: all-atom structure validation for macromolecular crystallography. *Acta Crystallogr D Biol Crystallogr* **66**, 12-21 (2010).
65. E. F. Pettersen *et al.*, UCSF ChimeraX: Structure visualization for researchers, educators, and developers. *Protein Sci* **30**, 70-82 (2021).
66. W. Chang, C. Tu, T. H. Chen, D. Bikle, D. Shoback, The extracellular calcium-sensing receptor (CaSR) is a critical modulator of skeletal development. *Sci Signal* **1**, ra1 (2008).

67. W. Chang *et al.*, PTH hypersecretion triggered by a GABA(B1) and Ca(2+)-sensing receptor heterocomplex in hyperparathyroidism. *Nat Metab* **2**, 243-255 (2020).
